# Unrestrained growth of correctly oriented microtubules instructs axonal microtubule orientation

**DOI:** 10.1101/2021.06.03.446862

**Authors:** Maximilian AH Jakobs, Assaf Zemel, Kristian Franze

## Abstract

In many eukaryotic cells, directed molecular transport occurs along microtubules. Within neuronal axons, transport over vast distances particularly relies on uniformly oriented microtubules, whose +-ends point towards the distal axon tip (+end out). However, axonal microtubules initially have mixed orientations, and how they orient during development is not yet fully understood. Using live imaging of primary *Drosophila melanogaster* neurons and physical modelling, we found that +end out microtubules are less likely to undergo catastrophe near the advancing axon tip, leading to their persistent long-term growth. In contrast, oppositely oriented microtubules remain short. Using chemical and physical perturbations of microtubule growth and genetic perturbations of the anti -catastrophe factor p150, which was enriched in the distal axon tip, we confirmed that the enhanced growth of +end out microtubules is critical for achieving uniform microtubule orientation. Computer simulations of axon development mimicking the enhanced +end out microtubule growth identified here along with previously proposed mechanisms correctly predicted the long-term evolution of axonal microtubule orientation as found in our experiments, highlighting the importance of the reduced catastrophe rate of +end out microtubules near the advancing axon tip in establishing uniform microtubule polarity. Our study thus leads to a holistic explanation of how axonal microtubules orient uniformly, a prerequisite for efficient long-range transport essential for neuronal functioning.

## Introduction

Symmetry breaking is critical for many biological systems. An organism starts of as a single round cell that divides and differentiates into many cells, tissues and organ systems. The neuron, with its branched dendrites and sometimes exceedingly long axon, is one of the least symmetric cells found in animals. Axons connect neurons with distant targets and thus enable long-distance signal transmission throughout the body at high speed.

The enormous length of axons, which can extend over several meters in some vertebrate species, poses substantial logistical challenges. RNA, proteins, and organelles originating in the cell body need to be actively transported down the axon. Transport occurs along MTs, which are long, polarized polymers that undergo stochastic cycles of growth and shrinkage **(Figure 1A)**. Motor proteins transport cargo either towards a MT’s dynamic (i.e., growing or shrinking) +end, or the stabilized -end.

**Figure 1:**
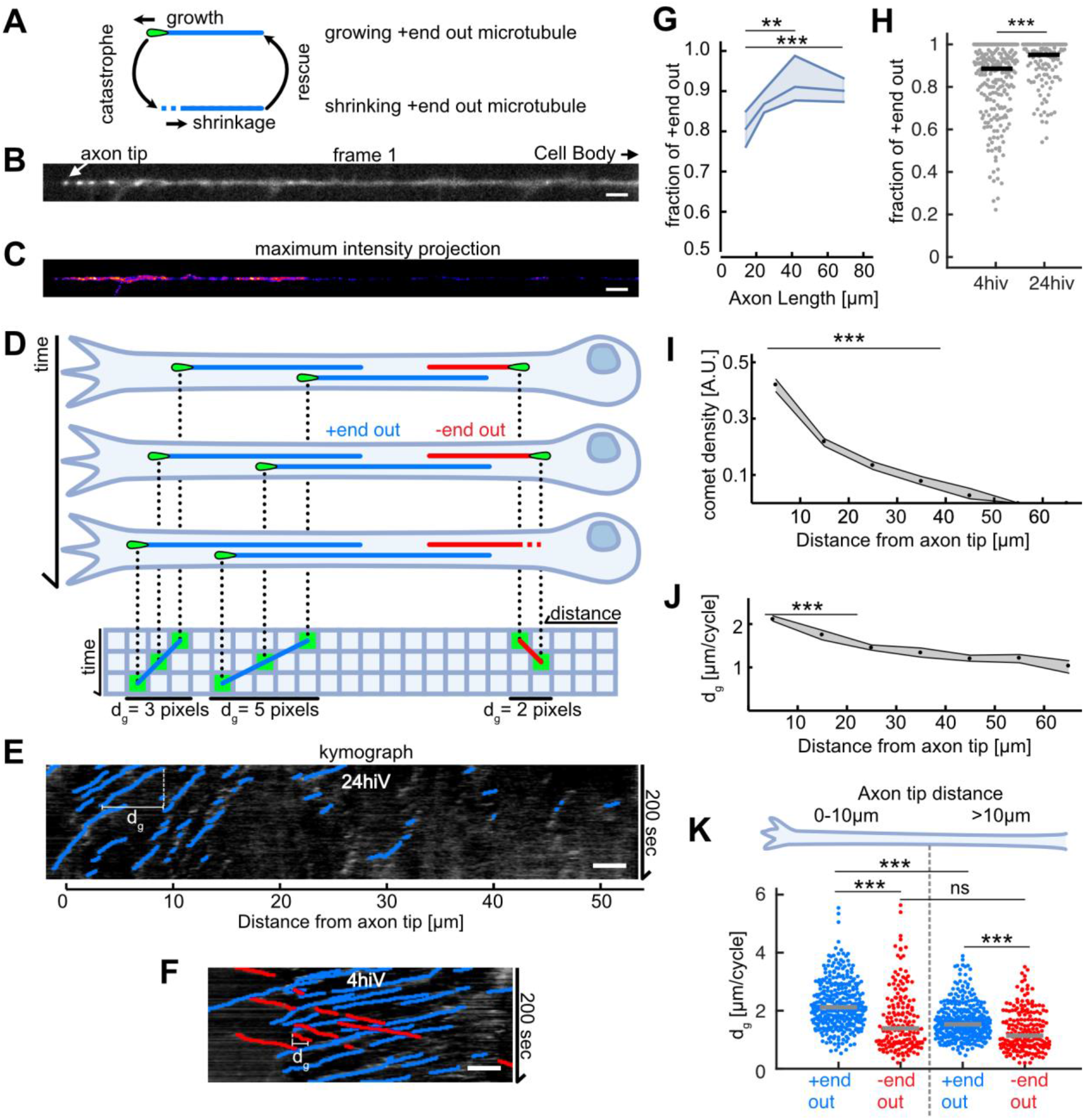
Axonal MT orientation increases over time and MT growth is enhanced at axon tips. **(A)** Schematic depicting the MT growth and shrinkage cycle. MTs grow until they undergo a catastrophe, which initiates MT shrinkage, and they start growing again after a rescue event. During growth (but not during shrinkage), EB1 localizes to MT tips. **(B)** First frame of a live cell imaging movie of axonal EB1-GFP dynamics. Bright dots represent individual EB1-GFP puncta, which label growing MT +ends. **(C)** Maximum intensity projection of a 200-second-long movie depicting EB1-GFP dynamics in a *D melanogaster* axon. EB1-GFP density is increased towards the tip. **(D)** Schematic showing how EB1-GFP live imaging movies were visualized and analysed using kymographs. The growing tips of +end out MTs (blue) and -end out MTs (red) were fluorescently labelled with EB1-GFP (green tear drop shaped ‘comets’). The same axon is shown at three different time points; MTs grow at their +end, where EB1-GFP is located. The axonal intensity profiles of all time points are plotted underneath each other, resulting in a space-time grid called ‘kymograph’. Connecting puncta between consecutive kymograph lines with blue/red lines yields the overall displacement *d_g_* for individual MT growth events. Note that the red -end out MT stops growing in the 2^nd^ frame and shrinks in the 3^rd^ frame **(E)** Kymograph of an axon 24 hours post plating showing EB1-GFP dynamics analysed with *KymoButler* (Jakobs et al., 2019). Lines with a positive slope (blue, left to right upwards) are MTs growing with their +end towards the axon tip, lines with a negative slope (red, left to right downwards) are MTs growing away from the tip. Horizontal bars indicate the growth lengths (dg) for individual MT growth cycles. **(F)** Kymographs of axonal processes expressing EB1-GFP analysed with *KymoButler* 4 hours post-plating **(G)** MT orientation as a function of axon length. Longer axons exhibit a more pronounced +end out MT orientation (p < 10^-5^, Kruskal Wallis test, **p < 0.01, ***p < 0.001 for pairwise comparisons, Dunn Sidak post hoc test). **(H)** MT orientation at 4 h.i.v and 24 h.i.v (*hours in vitro*). MT orientation increased with time (p < 10^-4^, Wilcoxon ranksum test). **(I)** EB1-GFP comet density as a function of the distance from the axon tip. Most MT polymerization occurred near the advancing axon tip. (N Axons = 353, 20 biological replicates from 20 different experiment days, p < 10^-20^, Kruskal Wallis test, p < 10^-7^ for pairwise comparisons between bin 1 to 2,3, or 4, Dunn-Sidak post hoc test). Shown are median ± 95% confidence interval. **(J)** Added length per MT growth cycle *d_g_* as a function of distance from the axon tip. MTs grew longer in the vicinity of the axon tip (p < 10^-20^, Kruskal Wallis test, p < 10^-7^ for pairwise comparisons of either bin 1 or 2 with any other bin, Dunn-Sidak post hoc test). **(K)** *d_g_* for +end out (blue) and -end out (red) MTs grouped for growth in the distalmost 10 μm of the axon tip, and further away than 10 μm from the axon tip. Each dot represents the average of one axon in the respective region, grey lines indicate median values. With *d_g_* 2.11 [2.04, 2.16] μm / cycle (bootstrapped median [95% confidence interval]), +end out MTs near the axon tip grew significantly longer than -end out MTs (*d_g_* =1.39 [1.27, 1.50] μm/cycle) and MTs located further away from the tip (*d_g_* = 1.53 [1.47, 1.59] μm / cycle, +end out, *d_g_* = 1.16 [1.03, 1.33] μm / cycle, -end out) (N = 346 (+end out close to tip), 343 (+end out away from tip), 177 (-end out close to tip), 194 (-end out away from tip) axons), 20 biological replicates; p < 10^-30^, Kruskal Wallis test followed by Dunn-Sidak post hoc test; *** p < 10^-4^). Scale bars: 3μm.

In immature axons, MT orientation is mixed, with 50-80% of all MTs pointing with their +end out (del Castillo et al., 2015; Yau et al., 2016a). During early neuronal development, the fraction of +end out axonal MTs increases (del Castillo et al., 2015; Yau et al., 2016a). In mature axons, ~95% of all MTs point in the same direction (+end out) (Baas et al., 1989; Heidemann et al., 1981), enabling polarized transport (Millecamps and Julien, 2013). Deficits in polarized transport have been associated with human neurodegenerative diseases, such as Alzheimer’s and Parkinson’s disease (Millecamps and Julien, 2013). Despite the importance of polarized transport in neuronal axons, the mechanisms that establish and maintain MT orientation are still not fully understood (Baas and Lin, 2011; Conde and Cáceres, 2009; Kapitein and Hoogenraad, 2011).

MTs in post-mitotic neurons are not attached to the centrosome (Kuijpers and Hoogenraad, 2011). Nucleation of new MTs occurs from MT organising centres (MTOCs) such as somatic Golgi (Mukherjee et al., 2011) through elongation of severed pieces (Yu et al., 2008) or *de novo* polymerization alongside existing MTs (Nguyen et al., 2014; Sánchez-Huertas et al., 2016). These newly formed MTs often orient in the same direction as existing ones, enforcing any pre-existing orientation bias (Mattie et al., 2010; Mukherjee et al., 2020). Pre-existing biases are furthermore enhanced by selective stabilisation of MTs through TRIM46-mediated parallel bundling (van Beuningen et al., 2015). However, without pre-existing biases these mechanisms by themselves cannot explain the robust +end out orientation of MTs in mature axons.

Furthermore, in axons, short MTs pointing with their -end away from the cell body can be transported towards the cell body (i.e., away from the tip) by cytoplasmic dynein (del Castillo et al., 2015; Rao et al., 2017), thus potentially clearing the axon of -end out MTs. To test whether this mechanism is sufficient to establish uniform microtubule orientation in axons, we previously designed computer simulations of dynein-mediated MT sliding in neurons. However, while MTs in the distal axon were oriented mostly with their +end away from the cell body {Jakobs:2015hk, Jakobs:2020jg}, our simulations failed to explain the longer-term +end out orientation of MTs in the proximal axon and the gradual establishment of a uniform +end out MT orientation throughout the axon seen in experiments ({Yau:2016jv}). Thus, our simulations suggested that additional mechanisms are needed to establish the uniform +end out orientation of MTs in neuronal axons.

Here, we investigated MT growth behaviours along *Drosophila melanogaster* axons and discovered a growth bias of +end out MTs near the advancing axon tip, which depended on the presence of MT anti-catastrophe protein gradients in the distal axon. A stochastic model of MT dynamics suggested that this growth bias leads to unbounded growth of these MTs, while -end out MTs remain short. Experiments and computer simulations confirmed that the observed selective growth bias is critical for uniform axonal MT orientation, leading to a model integrating known mechanisms with the decreased +end out MT catastrophe rate discovered here to explain how uniform MT orientation is achieved in developing neuronal axons.

## Results

### Correctly oriented MTs add more length per growth cycle

To investigate how MT orientation in neuronal axons becomes biased, we first cultured acutely dissociated neurons from the *Drosophila melanogaster* larval CNS (Egger et al., 2013) and quantified MT growth in axons. We used *Drosophila* lines expressing the fusion protein EB1-GFP, which labels growing MT +ends with bright ‘comets’ **(Figure 1A-C**) (Sanchez-Soriano et al., 2010; Stepanova et al., 2003). The distance over which a comet moves in the axon is equal to the overall length *d_g_* that is added to a MT between the start of its growth cycle and the catastrophe leading to MT shrinkage **(Figure 1A)**. The direction of growth reveals whether a MT is oriented with its +end away from (+end out) or towards (-end out) the cell body. Time-lapse movies of EB1-GFP comets were converted into kymographs and analysed using *KymoButler* (Jakobs et al., 2019) **(Figure 1D-F**).

The fraction of +end out MTs increased over time and with increasing axonal length **(Figure 1G-H**), confirming that MT orientation increases during development (del Castillo et al., 2015; Yau et al., 2016a). Most MT growth events (~66%) were found within the first 20 μm from the advancing axon tip **(Figure 1A, J**). MT growth lengths per cycle, *d_g_*, were significantly higher near the axon tip compared to further away from it **(Figure 1K**). Furthermore, +end out MTs added significantly more length per growth cycle than -end out MTs, with the highest difference between +end out and -end out MTs (~0.5 μm/cycle) found within the first 10 μm from the axon tip (*d_g_* (+end out)= 2.11 [2.04, 2.16] μm / cycle & *d_g_* (-end out)= 1.39 [1.27, 1.50] μm, bootstrapped median [95% confidence interval]) **(Figure 1L**).

Increases in MT growth lengths during a polymerisation cycle *d_g_* = *v_g_/f_g_* could either arise from an increased polymerization velocity *v_g_* or a decreased catastrophe frequency *f_g_* (or both). In *drosophila* neurons, MT growth velocities *v_g_* were similar in all MTs irrespective of their orientation and position within the axon (~5 μm / min; Error! Reference source not found.**C**). Additionally, EB1-GFP, which affects microtubule polymerisation velocities, was not enriched at the axon tip (Error! Reference source not found.**E**), and EB1-GFP comet lengths, which have previously been linked to microtubule polymerisation speeds (Roostalu et al., 2020), did not show significant variations along the axon (Error! Reference source not found.**F**). However, catastrophe rates *f_g_* = 1/*t_g_* were significantly lower (and thus polymerization times *t_g_* significantly longer) in +end out MTs near the axon tip (*f_g+_*~4 * 10^−2^*sec*^−1^ *vs. f_g−_*~ 6 * 10^−2^*sec*^−1^; Error! Reference source not found.**B**), suggesting that these MTs grew longer because of a decrease in their catastrophe frequencies.

Overall, the orientation of MTs that added more length per growth cycle (i.e., +end out MTs) becomes the dominant MT orientation in developing axons, indicating a possible link between increased +end out MT growth and the fraction of +end out MTs in axons.

### Enhanced growth of +end out MTs leads to unbounded growth

On long time scales, differences in MT growth length per growth cycle accumulate and thus affect the average MT length *l_MT_*. To estimate whether the rather small differences in *d_g_* of ~0.5 μm/cycle might lead to biologically meaningful differences in the average expected MT lengths between +end out and -end out oriented MTs, we used a 2-state Master equation model of MT growth and shrinkage (see (Dogterom and Leibler, 1993) & supplemental methods for details). The model distinguishes two regimes **(Figure 2A**):

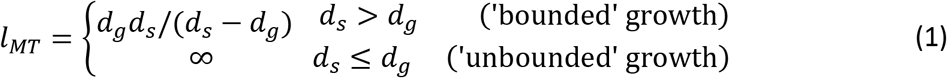

where *d_s_* = lost length per shrinkage cycle.

**Figure 2:**
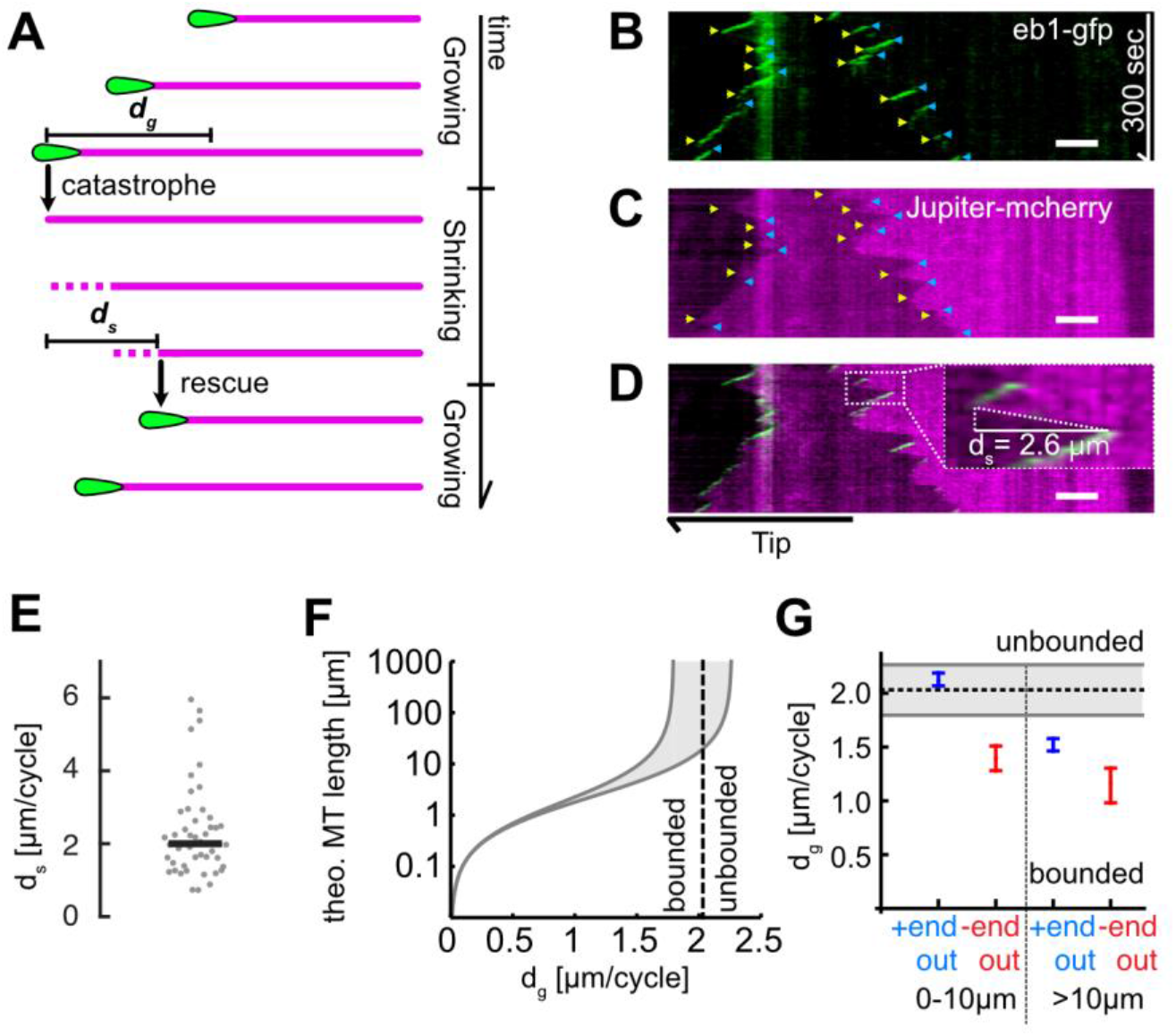
MT length depends on added length per growth cycle. (**A)** Schematic highlighting the assumptions of our two-state master equation model. MTs were assumed to occupy either a growing or shrinking state. During a growth cycle, the average MT length increases by *d_g_*, during a shrinkage cycle, the MT length decreases by *d_s_*. Additionally, MTs were able to stochastically switch between the two states as shown in **Figure 1A. (B-D)** Kymographs from a *D. melanogaster* axon that expressed **(B)** EB1-GFP (green) and **(C)** Jupiter-mCherry, a tubulin label (magenta). Individual MT shrinkage events, visible as **(C)** fluorescent edges and **(D)** dashed white lines in the kymograph, yielded MT shrinkage lengths per cycle *d_s_*. Yellow and blue arrow heads in **B** & **C** indicate start and end points of an individual shrinkage event, and the inset in **D** highlights an individual shrinkage event. Scale bars: 3μm. **(E)** Average *d_s_* values for N=47 axons (3 biological replicates; median: 2.03 [1.80, 2.26] μm (bootstrapped median [95% confidence]). **(F)** Plot of the estimated overall MT length *l_MT_* as a function of *d_g_*. The two solid black curves indicate the lower and upper bounds of the average MT lengths for a given *d_g_* with *d_s_* = 1.80 or 2.26 μm. One can separate two regimes, “unbounded” and “bounded” growth, separated by the dashed grey line. **(G)** Plot of *d_g_* as a function of MT orientation and localisation showing median and 95% confidence intervals. +end out MTs close to the tip were considerably more likely to exhibit unbounded growth than +end out MTs further away from the tip and -end out MTs.

When *d_s_* ≤ *d_g_*, the average length added to the MT exceeds the average shrinkage length per cycle so that an MT will exhibit net growth and elongate if physically possible in its confined environment (called ‘unbounded’ growth). For *d_s_* > *d_g_*, however, growth is ‘bounded’, and average MT lengths follow an exponential distribution with a mean of *d_s_d_g_*/(*d_s_-d_g_*). In practice, this means that MTs with *d_g_ ≥ d_s_* would grow until encountering a physical barrier (for example the distal end of the axon tip), while MTs with *d_s_* > *d_g_* remain finite (for example, approximately 2 μm with *d_s_* = 2 μm and *d_g_* = 1.5 μm).

We determined the MT shrinkage per cycle *d_s_* by co-expressing a Jupiter-mCherry fusion protein (a tubulin marker) together with EB1-GFP in *D. melanogaster* axons. MTs stopped growing when the GFP signal disappeared from their +end, indicating a catastrophe or pause event. Subsequent MT shrinkage was visualized by simultaneously imaging tubulin (Jupiter-mCherry) and quantified by tracing tubulin edges resulting from the shrinkage in the dual colour kymographs **(Figure 2B-D**). Axonal MT shrinkage lengths were *d_s_* = 2.03 [1.80, 2.26] μm / cycle (bootstrapped median [95% confidence interval], **Figure 2E**).

With this value for *d_s_*, our model predicted the divergence of *l_MT_* at *d_g_* = 2.03 μm, at the lower end of the measured 95% confidence interval for the median *d_g_* = [2.04, 2.16] μm / cycle of +end out MTs near the axon tip but well above the 95% confidence interval for the median *d_g_* = [1.39, 1.48] μm / cycle of all other MTs **(Figure 2F).** The measured values of *d_g_ and d_s_* hence suggested that +end out-oriented MTs exhibit mostly unbounded growth within 10 μm from the axon tip while -end out MTs within that range and any MT further away from the tip do not **(Figure 2F)**. The decreased catastrophe frequency of +end out MTs near the axon tip (Error! Reference source not found.**A-C**) implied a higher chance of survival for +end out MTs while leaving -end out MTs labile, thereby establishing a bias for +end out microtubule orientation.

### Enhanced growth of +end out MTs is required for uniform +end out MT orientation

To test if enhanced MT growth is indeed involved in biasing MT orientation, we first chemically decreased MT growth using Nocodazole, a drug that disrupts MT polymerization. Nocodazole treatment led to decreased MT growth velocities *v_g_* (**Error! Reference source not found.**). Alternatively, we also physically decreased MT growth by increasing the osmolarity of the cell culture medium through addition of NaCl (BRAY et al., 1991; Molines et al., 2020). Here, MT catastrophe rates *f_g_* were significantly increased (**Error! Reference source not found.**).

Both treatments led to significantly decreased +end out MT growth *d_g_ = v_g_/f_g_* at the axon tip (< 10 μm), with *d_g_* < 1.71 μm / cycle < *d_s_* (upper 95% confidence bound of median) **(Figure 3A-E**). Our model predicted that this change in *d_g_* should lead to a switch from previously unbounded to bounded growth of +end out MTs near the axon tip **(Figure 3D**), thus reducing the bias towards +end out MTs and decreasing MT orientation. In agreement, MTs within treated axons were overall significantly less uniformly oriented **(Figure 3D-E**), confirming an important role of the enhanced growth lengths per cycle of +end out MTs in establishing axonal MT orientation.

**Figure 3:**
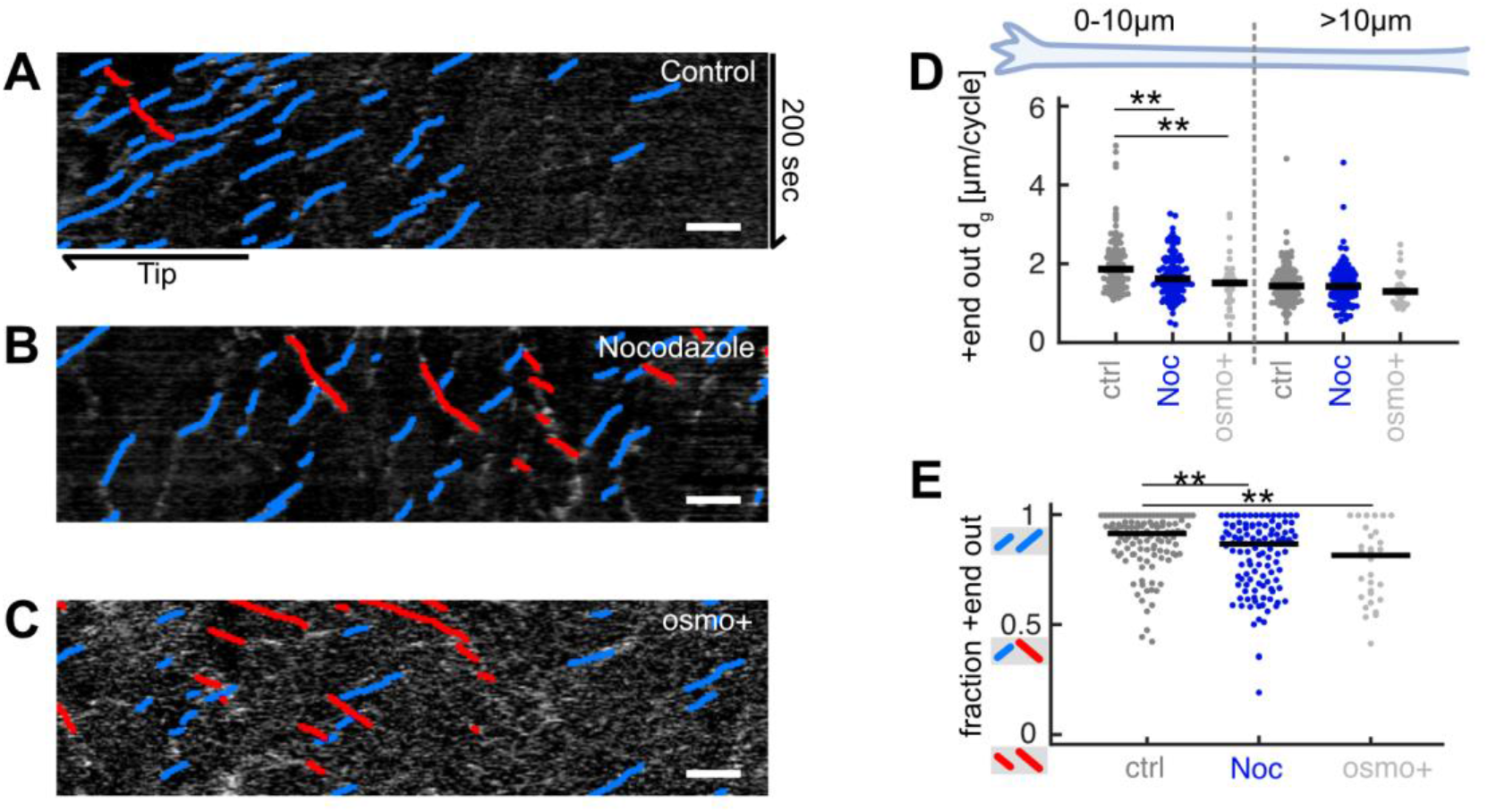
Decreasing MT growth leads to decreased axonal MT orientation. **(A-C)** Representative kymographs analysed with *KymoButler* (Jakobs et al., 2019) from axonal processes treated with (**A**) 0.025% DMSO (control) for 8h, **(B)** 5uM Nocodazole for 8h, **(C)** medium with increased osmolarity (“osmo+”) for 22h. Growth of +end out MTs is shown as blue lines, -end MTs are red. Scale bars = 3μm. **(D)** Added MT lengths per growth cycle *d_g_* of +end out MTs at the distalmost 10μm from the axon tip and further away for control (N = 107 axons from 5 biological replicates), nocodazole-treated axons (N = 116 axons from 3 biological replicates), and axons cultured in osmo+ medium (N = 30, 2 biological replicates). At the axon tip, MT lengths increased significantly less per growth cycle in axons treated with Nocodazole or osmo+ media than controls (p < 10^-4^, Kruskal Wallis test, ** p < 0.01 for pairwise comparisons, Dunn-Sidak post hoc test). **(E)** The fraction of +end out MTs in the different groups. MT orientation was calculated by counting all MTs that grew away from the cell body (blue lines in kymographs) and dividing them by all growing MTs (blue and red) along the whole axons. This way, a kymograph with only blue lines gives a value of 1 while an equal number of blue and red lines yields a value of 0.5. MTs in axons treated with Nocodazole or osmo+ media were significantly less uniformly oriented than those in the control group, i.e., they contained a larger fraction of MTs pointing with their +ends toward the cell body (red lines in **(A-C)**) (p < 10^-4^, Kruskal Wallis test, ** p < 10^-2^ for pairwise comparisons, Dunn-Sidak post hoc test).

### p150 protein gradient in axon tips promotes +end out MT stabilisation

However, why do +end out MTs grow longer in the vicinity of the axon tip? Local gradients of MT growth-promoting factors could lead to an increase in +end MT growth in that region. Axon tips contain a multitude of different proteins and are highly compartmentalised (Lowery and Van Vactor, 2009). Locally enriched MT growth-promoting factors (of which there are many in the axon tip (Voelzmann et al., 2016)), include MT stabilizing proteins, such as p150 (Lazarus et al., 2013; Moughamian and Holzbaur, 2012), CRMP-2 (Fukata et al., 2002; Inagaki et al., 2001), and TRIM46 (Rao et al., 2017; van Beuningen et al., 2015), as well as free tubulin and others (Eng et al., 1999).

The MT stabilizing protein p150, for example, affects MT catastrophe rates *f_g_* and nucleation rates but not growth velocities *v_g_* (Lazarus et al., 2013). As we observed decreased MT catastrophe rates but constant growth velocities of +end out MTs towards axon tips (**Error! Reference source not found.B, C**), we hypothesised that p150 is one of the key proteins involved. *Drosophila* has a p150 homologue which, similar as in murine neurons (Moughamian and Holzbaur, 2012), we found to be enriched in axonal but not in dendritic tips **(Figure 4A, C** and **Error! Reference source not found.**). Accordingly, +end out MT growth lengths per cycle *d_g_* and catastrophe rates *f_g_* were significantly higher in axons than in dendritic processes (Error! Reference source not found.**E,G**), where *d_g_* = 1.42 [1.37, 1.39] μm / cycle remained smaller than *d_s_* = 2.03 [1.80, 2.26] μm / cycle. In agreement with our model, which predicted unbounded growth of +end out MTs in axons but bounded growth in dendritic processes (**Figure 2**), dendritic processes exhibited mixed (50% +end out) MT orientations (Error! Reference source not found.**D**), corroborating a link between MT stabilizing protein gradients in the tip of axons, reduced MT catastrophe rates, unbounded growth, and overall MT orientation.

**Figure 4:**
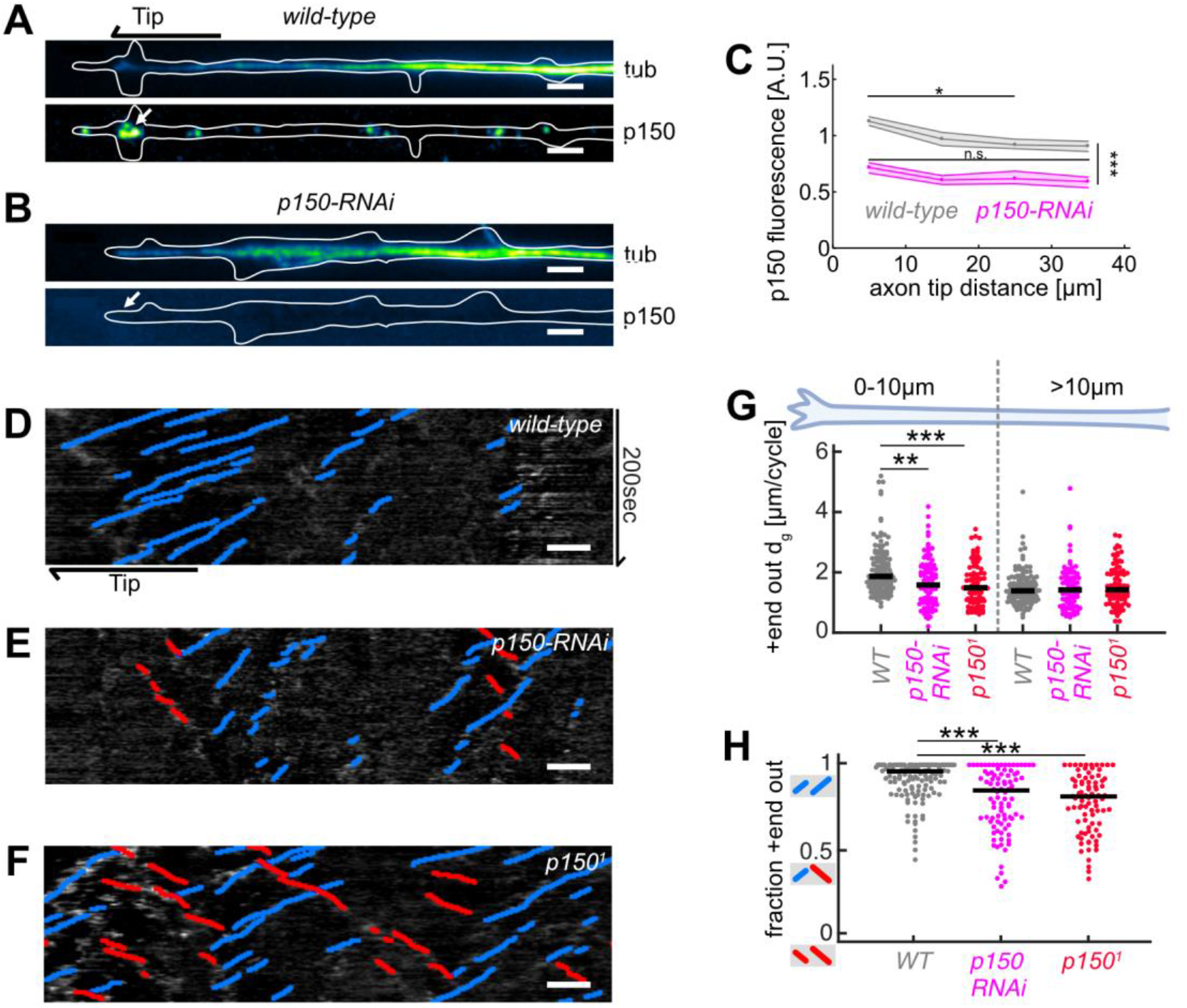
Abrogation of p150 function decreases MT growth and axonal MT orientation. **(A-B)** Tubulin (top) and normalized p150 (bottom) immunostaining of cultured *D. melanogaster* larvae axonal processes of **(A)** controls and **(B)** neurons expressing *elav-gal4* UAS-driven *p150-RNAi*. Large p150 puncta were found clustered around the axon tip (arrow) in controls **(A)** but not in *p150-RNAi* axons **(B)**. Scale bars = 2μm **(C)** Normalized p150 fluorescence intensity as a function of distance from the axon tip for *wild-type* axons (N = 83, 2 biological replicates) and *p150-RNAi* axons (N = 111, 2 biological replicates). Lines represent median ± 95% confidence intervals for *wild-type* (grey) and *p150-RNAi* (magenta). P150 fluorescence intensities changed along the axon (p < 10^-70^; Kruskal Wallis test). In *wild-type* axons, p150 was enriched at the axon tip (* p < 0.05 between bin 1 and bin 3 or 4; pairwise comparisons with Dunn-Sidak post hoc test), but not in *p150-RNAi* expressing axons (p > 0.05 for all pairwise comparisons). Overall, p150 expression levels were diminished in *p150-RNAi* axons compared to *wild-type* (*** p < 10^-7^ for any pairwise comparison between conditions). **(D-F)** *KymoButler* output for kymographs of EB1-GFP expressed in (**D**) a *wild-type* axon, **(E)** an axon expressing p150-RNAi, and **(F)** an axon in a *p150^1^/+* mutant background. Scale bars = 3 μm. Blue/red lines represent MTs with +/-end out orientation, respectively. **(G)** +end out MT added length per growth cycle *d_g_* for *wild-type* (N = 85, 9 biological replicates), *p150-RNAi* (N = 34, 3 biological replicates), and *p150^1^/+* (N = 83, 6 biological replicates). At the axon tip, MT growth lengths were significantly decreased in both *p150-RNAi* and *p150^1^* conditions compared to controls (p < 10^-9^, Kruskal Wallis test, ***p < 0.001, *p < 0.05, Dunn-Sidak post hoc test). **(H)** MT orientation along the whole axon for *wild-type, p150-RNAi*, and *p150^1^/+*. MTs were less uniformly oriented in both *p150-RNAi* and *p150^1^* axons (p < 10^-9^, Kruskal Wallis test, *** p < 10^-5^ for pairwise comparisons with Dunn-Sidak post hoc test). Overall, axonal MT orientation was decreased after chemical, physical, and genetic perturbations of MT growth.

We next tested whether the observed p150 gradient is strong enough to lead to different MT growth lengths (via modulation of catastrophe rates, *d_g_ ~ 1/f_g_*) for +end out and -end out MTs within 10 μm from the axon tip, the region where we observed the largest differences in *f_g_* (**Error! Reference source not found.C**). We assumed that MT growth lengths per cycle at a distance *x* from the axon tip can be described by a power law function *d_g_* (*x*)= *A*\**p150*(*x*)*^α^* of the p150 fluorescence intensity profile. A simultaneous fit to both +end out and -end out MTs demonstrated that the observed gradient is, in theory, indeed strong enough to cause different growth behaviours for +end out and -end out MTs in axonal tips (Error! Reference source not found.).

To test this prediction further, we assessed MT dynamics in *wild-type, p150-RNAi* expressing neurons, and in neurons from *p150^1^/+* mutant flies (Figure 4). *p150^1^* (also known as *Gl^1^*) mutants express a truncated p150-RNA transcript, which results in a dominant negative phenotype (Plough and Ives, 1935). Expressing *p150-RNAi* led to decreased p150 protein in axons **(Figure 4A-C)**. Both the expression of *p150-RNAi* and of dominant negative *p150^1^/+* led to a significant increase in +end out MT catastrophe rates (**Error! Reference source not found.**) and thereby to a decrease (14%-20%) in +end out MT growth within 10 μm from the axon tip **(Figure 4G**). In agreement with our model, the overall axonal MT orientation was significantly decreased in both *p150-RNAi* and *p150^1^/+* axons compared to controls **(Figure 4H**).

p150 is known to interact with dynein (Karki and Holzbaur, 1995). Expressing an RNAi against dynein heavy chain also led to decreased +end out MT growth at the axon tip (25%) and decreased MT organisation, reproducing the results of p150 removal (Error! Reference source not found.). Together, these data indicated that MT stabilising or growth-promoting protein gradients at the axon tip do indeed have an important role in regulating the overall orientation of the axonal MT network.

### Kinesin 1 is required to establish p150 gradient

Previous work showed that p150 accumulation at axon tips depends on the activity of the MT-specific molecular motor protein kinesin 1 (Moughamian and Holzbaur, 2012; Twelvetrees et al., 2016), which preferentially enters axons over dendrites (Tas et al., 2017). Accordingly, disruption of kinesin 1 function with an RNAi treatment led to a 25% decrease in p150 fluorescence within 10 μm from the axon tip (Error! Reference source not found.**A-D**). Again, the absence of a p150 gradient in these neurons led to a significant increase in MT catastrophe rates and decreased +end out MT growth near the axon tip, and hence to an overall decrease in axonal MT orientation (Error! Reference source not found.**E-I**), suggesting that gradients of MT-stabilising proteins at the tips of developing axons are critical for biasing axonal MT orientations.

### Uniform axonal MT orientation is established through a combination of MT sliding, templating, and unbounded growth

Our experiments suggested that a gradient of a MT growth-promoting factor localised at the axon tip is required for establishing uniform +end out MT orientation in axons. To investigate the significance of the locally biased microtubule growth in more detailed, we modelled the evolution of overall microtubule polarity along the axon using computer simulations. We extended our previously described simulations (Jakobs et al., 2020) to include two additional effects on axonal MTs: (1) enhanced stabilization of MTs at axon tips as found in this study, and (2) a local biasing of MT nucleation (MT templating) that could originate from augmin (Nguyen et al., 2014; Sánchez-Huertas et al., 2016) or TRIM46 (van Beuningen et al., 2015).

The simulation is detailed in the methods section and a schematic can be found in **Figure 5A**. Briefly, a cylindrical bundle of MTs is generated, and dynein motors are assumed to cross-link adjacent MTs. The orientation of each MT is randomly chosen. Force-velocity relationships are used to predict the exerted sliding velocities of the MTs, which are then used to calculate new MT positions iteratively. MTs were confined in the axon with a solid boundary at the axon tip and a semi-permeable boundary at the cell body. New MTs were added randomly along the bundle axis at a determined frequency. To mimic the effect of a concentration gradient of a growth-promoting protein (such as p150) at the axon tip, we assumed that the frequency of newly added MTs is sampled from an exponential distribution that peaks at the axon tip. To account for augmin or TRIM46-induced local biases of MT nucleation, we assumed that the orientation of an added MT is dictated by the mean MT-orientation in the region to which it is added.

**Figure 5:**
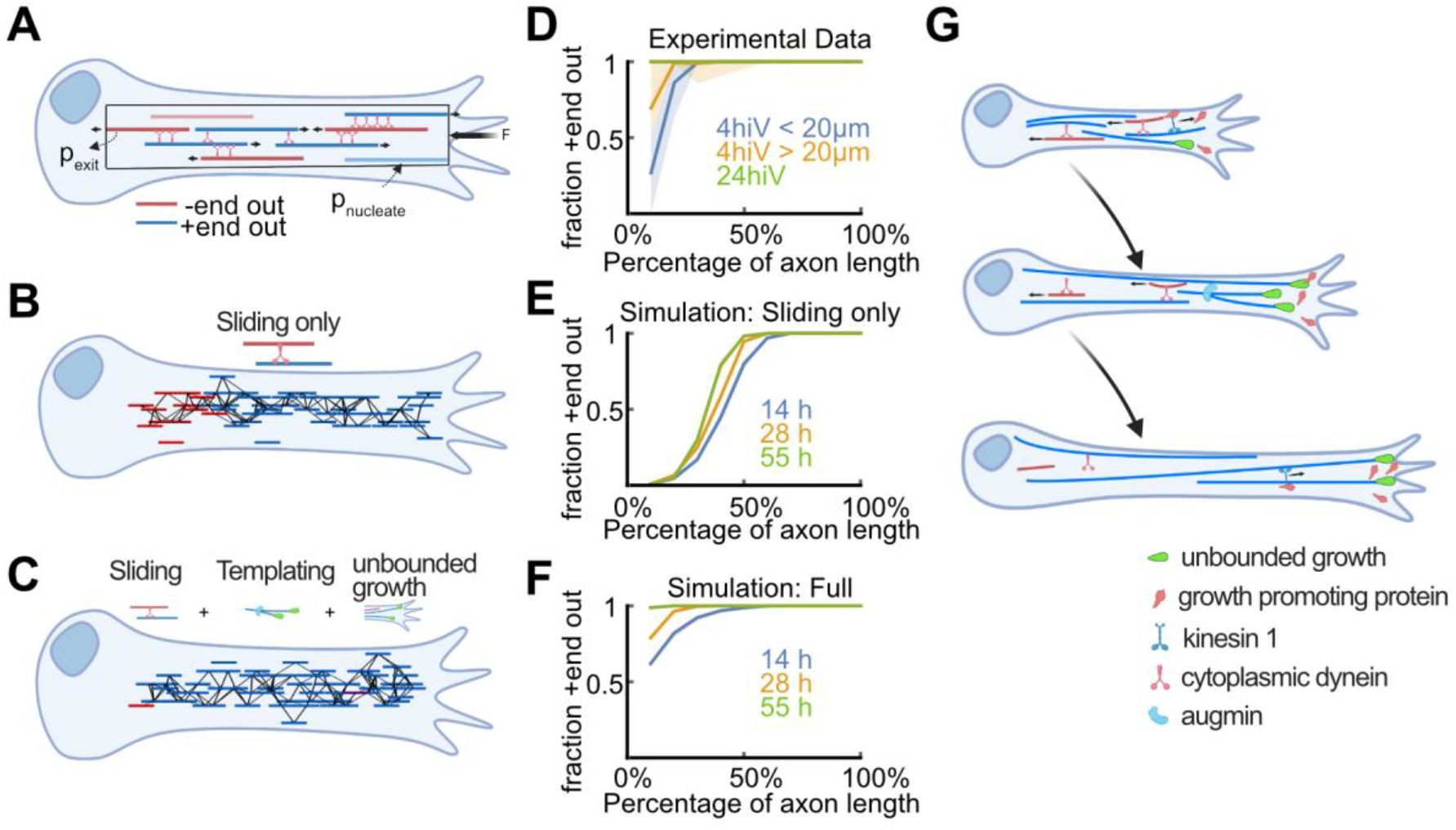
Biased stabilization of +end out MTs is required to establish uniform axonal MT orientation. **(A)** Schematic showing the MT sliding simulation and its relevant parameters. **(B-C)** Simulation results of MT dynamics. **(B)** Snapshot of a simulated axon with dynein-based sliding of MTs. -end out MTs accumulated within the proximal axon. **(C)** Snapshot of an axon simulated with sliding, augmin templating (new MTs were likely oriented into the same direction as their surrounding ones), and biased nucleation of +end out MTs at the tip. Much like in real axons, most MTs were oriented with their +end out throughout the axon. **(D)** Experimental profiles of MT orientation along the (normalized) axon length for axons that were cultured 4 hours in vitro (4hiV), separated into less than 20 μm or more than 20 μm long, and 24 hours in culture (24hiV). Lines represent bootstrapped medians and 95% confidence intervals. **(E-F)** Simulation profiles of MT orientation along the axon. **(E)** Profiles obtained from simulations with dynein-mediated sliding only at 3 different simulation timepoints. MT orientation was graded along the axon but, unlike in the experimental profiles shown in **(D)**, the proximal axon remained enriched with -end out MTs. **(F)** MT orientation profiles obtained from simulations with sliding, templating, and biased nucleation of +end out MTs at the axon tip. The observed gradual development of MT orientation along the axon is in excellent agreement with our experimental data **(D)**. **(G)** Summary of proposed mechanism for establishing MT orientation in axons. Red and blue lines represent -end out and +end out MTs, respectively. Green drop shapes indicate unbounded MT growth into the axon tip for +end out MTs. Kinesin 1 deposits MT growth promoting proteins, such as p150, at axon tips (Error! Reference source not found.), leading to local unbounded growth of +end out MTs. Augmin templating and cell body-directed sliding of -end out MTs further amplifies this bias. All three mechanisms together lead to a +end out MT cytoskeleton.

Simulations incorporating dynein-induced MT sliding but lacking mechanisms (1) and (2) mentioned above resulted in steadily growing axons whose MT orientation profile was graded along their length, being enriched with +end out MTs at the distal end and with –end out MTs at the proximal (cell body) end **(Figure 5A,B, E** and Error! Reference source not found.). The proximal domain of –end out MTs grew in proportion to the axon length, as –end out MTs from across the growing axon continually accumulated in that region despite their withdrawal into the cell body by dynein motors. Those simulations thus failed to reproduce the experimental observation of an enrichment of the proximal axon with +end out MTs over time, which eventually leads to uniformly oriented axonal MTs (Yau et al., 2016b) **(Figure 5D**).

Similarly, the increased growth of +end out microtubules or templating alone, or any combination of two out of these three mechanisms, were also insufficient for establishing high fractions of +end out MTs at the proximal axon as observed experimentally (Error! Reference source not found.).

However, when we integrated sliding, templating, and biased MT growth at the axon tip in the simulation (assuming that the likelihood of exhibiting unbounded growth corresponds to a successful nucleation event), MTs gradually oriented uniformly +end out across the entire length of the axon, recapitulating our experimental results **(Figure 5C, D, F** and Error! Reference source not found.). Thus, our data suggest that multiple mechanisms – including biased growth of +end out MTs near the axon tip identified in this study – need to work in unison to establish and maintain uniform axonal microtubule orientation.

## Discussion

We here found that MT growth is an important factor in the regulation of overall MT organisation in the axon. Our experiments and modelling suggest that an enrichment of MT stabilising / growth-promoting proteins at the advancing axon tip leads to a transition of MT growth from a bounded to an unbounded state for +end out MTs. This growth transition is important for establishing uniform +end out MT orientation from the cell body to the axon tip as found in mature axons. While previous studies suggested that MT dynamics are temporally and spatially constant during early axon formation (Seetapun and Odde, 2010), our results suggest that, at later stages of axon maturation, MT dynamics are heterogeneous (Figure 1).

Our chemical and physical perturbations of MT growth affected different aspects of MT dynamics. Nocodazole treatment decreased MT growth velocities while not affecting catastrophe frequencies; hyperosmotic solutions increased MT catastrophe frequencies but did not alter growth velocities (**Error! Reference source not found.**). The effect of the hyperosmotic solution might potentially arise from a decrease in the available space in the distal axon for MTs to polymerize into (Dogterom and Yurke, 1997; Franze, 2020). However, NaCl can be toxic for neurons, so that the effect on MT growth could also be due to a general stress response (Morland et al., 2016). Either way, both treatments led to a decrease of the microtubule length added during each growth cycle, *d_g_*, so that *d_g_ < d_s_*, switching +end out MT dynamics from unbounded to bounded growth, thus leading to a loss of uniform MT polarisation along the axon (Figure 3).

The axon tip is highly enriched with growth promoting factors such as p150 (Lazarus et al., 2013; Moughamian and Holzbaur, 2012), CRMP-2 (Fukata et al., 2002; Inagaki et al., 2001), TRIM46 (Rao et al., 2017; van Beuningen et al., 2015) and EB1 (Ma et al., 2004; Morrison et al., 2002). The MT stabilising protein p150, which we investigated here as an example, was concentrated at axon tips but was not enriched at dendritic tips (**Error! Reference source not found.).** Enhanced stabilisation of +end out MTs, leading to reduced catastrophe rates and thus unbounded growth essential for establishing uniform MT orientation, was only observed near axonal but not near dendritic tips (Error! Reference source not found.), confirming that differences in the localization of MT growth-promoting proteins correlate with differences in MT growth. Perturbations of p150 led to increased catastrophe rates and decreased +end out MT growth in the axon tip, and thus to decreased overall MT order in the axon (Figure 3).

While p150 is mainly known for of its role in the dynactin complex, which is an important cargo adapter protein complex for the molecular motor protein dynein (Gill et al., 1991), it also acts as a dynein-independent MT anti-catastrophe factor (Lazarus et al., 2013). Since p150 and dynein are functionally related, it is difficult to separate their individual contributions to stabilising +end out MT growth in the axon tip and cell body-directed sliding of -end out MTs along the axon. However, it remains unclear whether p150 is required for dynein-mediated MT sliding (Ahmad et al., 1998; Waterman-Storer et al., 1997), and the *Drosophila melanogaster* oocyte also contains a biased MT cytoskeleton whose orientation is, presumably, maintained by p150 (Nieuwburg et al., 2017). Hence, while p150 is unlikely to induce unbounded MT growth alone, it emerges as a key contributor to the establishment of MT orientation.

In addition to its contribution to setting up the p150 gradient in axon tips (Error! Reference source not found.), kinesin 1 is also thought to slide MTs with their -end leading (del Castillo et al., 2015) and to be involved in dynein function (Pilling et al., 2006; Rao et al., 2017). Kinesin knockdown could thus not only prevent the accumulation of MT growth-promoting factors at the axon tip but also decrease kinesin 1-mediated -end out MTs sliding into the distal axon (thereby increasing the fraction of +end out MTs) and/or prevent retrograde dynein-mediated sliding of - end out MTs (decreasing the fraction of +end out MTs). We observed that disruption of kinesin 1 function led to a decrease in the fraction of +end out MTs in the axon (Error! Reference source not found.), suggesting that kinesin 1 does not contribute to the overall MT orientation via direct sliding but that it rather affects MT orientation mainly via localising MT growth or nucleation-promoting proteins to axonal tips and/or via interactions with dynein-mediated sliding.

Finally, both kinesin 1 and p150/dynactin perturbations could potentially also affect MTOC localization in neurons. p150 was enriched at the tips of axonal but not of dendritic processes (Error! Reference source not found.). In *C. elegans* neurons, MTOCs may be located to the tips of dendritic processes (Liang et al., 2020). Removal of dynactin, which initiates cell body-directed transport from axonal tips (Moughamian and Holzbaur, 2012), could lead to an increased number of MTOCs also at axon tips, thereby promoting growth and nucleation of MTs. However, our results showed a decrease in MT growth dynamics at axon tips after dynactin removal (Figure 3), indicating that the observed decrease in MT orientation was mostly due to decreased rather than promoted growth of axonal MTs in the axon tip.

We propose the following model explaining the spontaneous establishment of MT orientation in developing axons. Growth-promoting proteins accumulate at the axon tip due to MT +end-directed transport by kinesin motors (Error! Reference source not found.). The resulting protein gradient leads to a local bias in MT growth (Figure 1, Figure 3), rendering +end out MT growth into the axon tip unbounded (Figure 2). In theory, a finite pool of tubulin at axon tips would further contribute to this bias in +end out MT growth by diminishing the amount of free tubulin available for MTs that polymerise towards the cell body. In contrast, short -end out MTs are more prone to depolymerization and/or transport away from the tip by dynein-mediated cell body-directed sliding (del Castillo et al., 2015; Rao et al., 2017), thus contributing to the orientation bias of MTs in the axon. The orientation bias is further enhanced by augmin-mediated templating or TRIM46-mediated parallel bundling of newly formed MTs to establish and maintain a fully organized MT cytoskeleton (see **Figure 5E** for a schematic summary). Together with cell process length-dependent MT accumulation (Seetapun and Odde, 2010), these mechanisms cooperate to build the polarized MT network that enables efficient long-range transport in neuronal axons. Future work will reveal whether other cellular systems use similar mechanisms to organize their cytoskeleton.

## Materials and Methods

### Key Resource Table

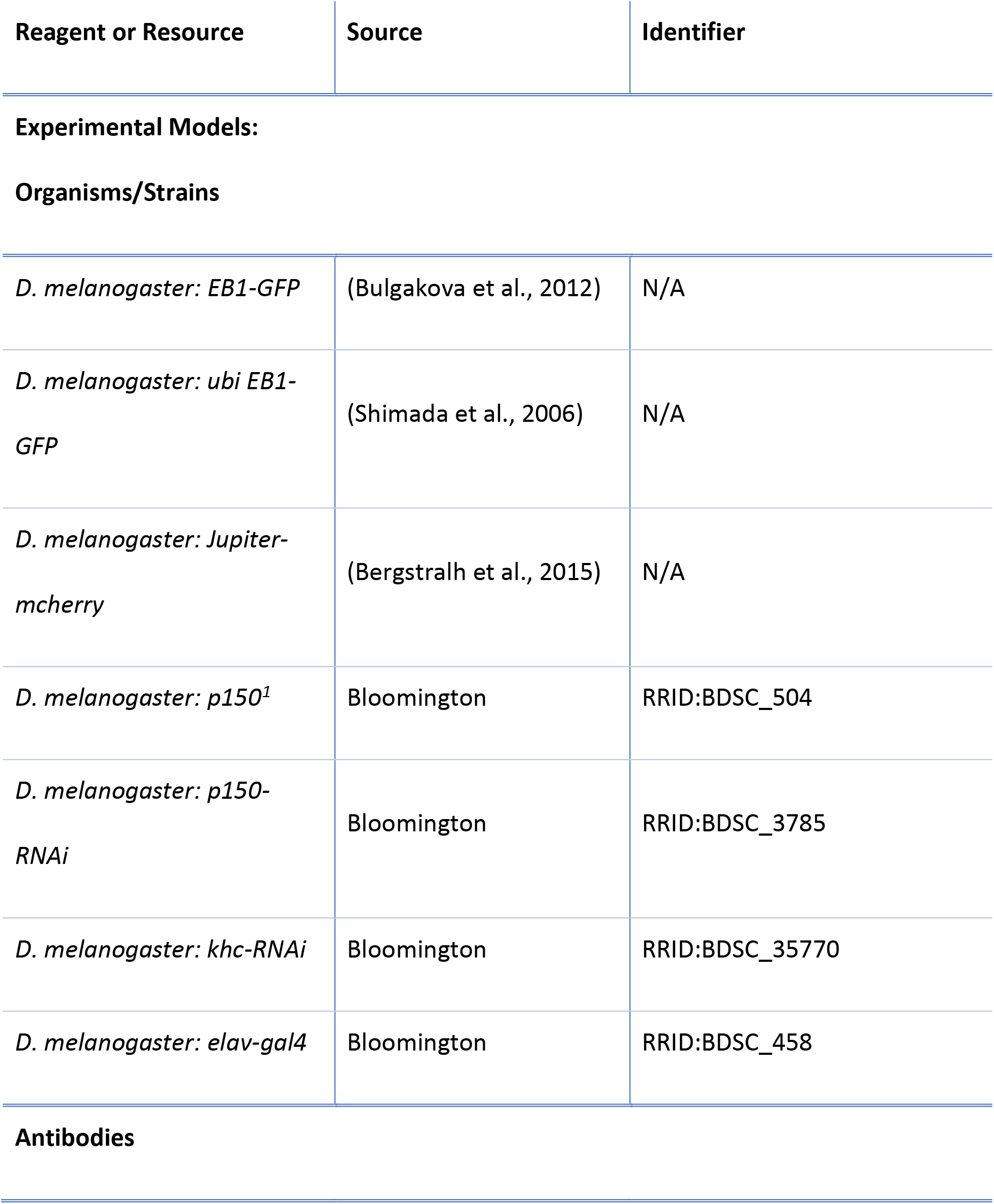
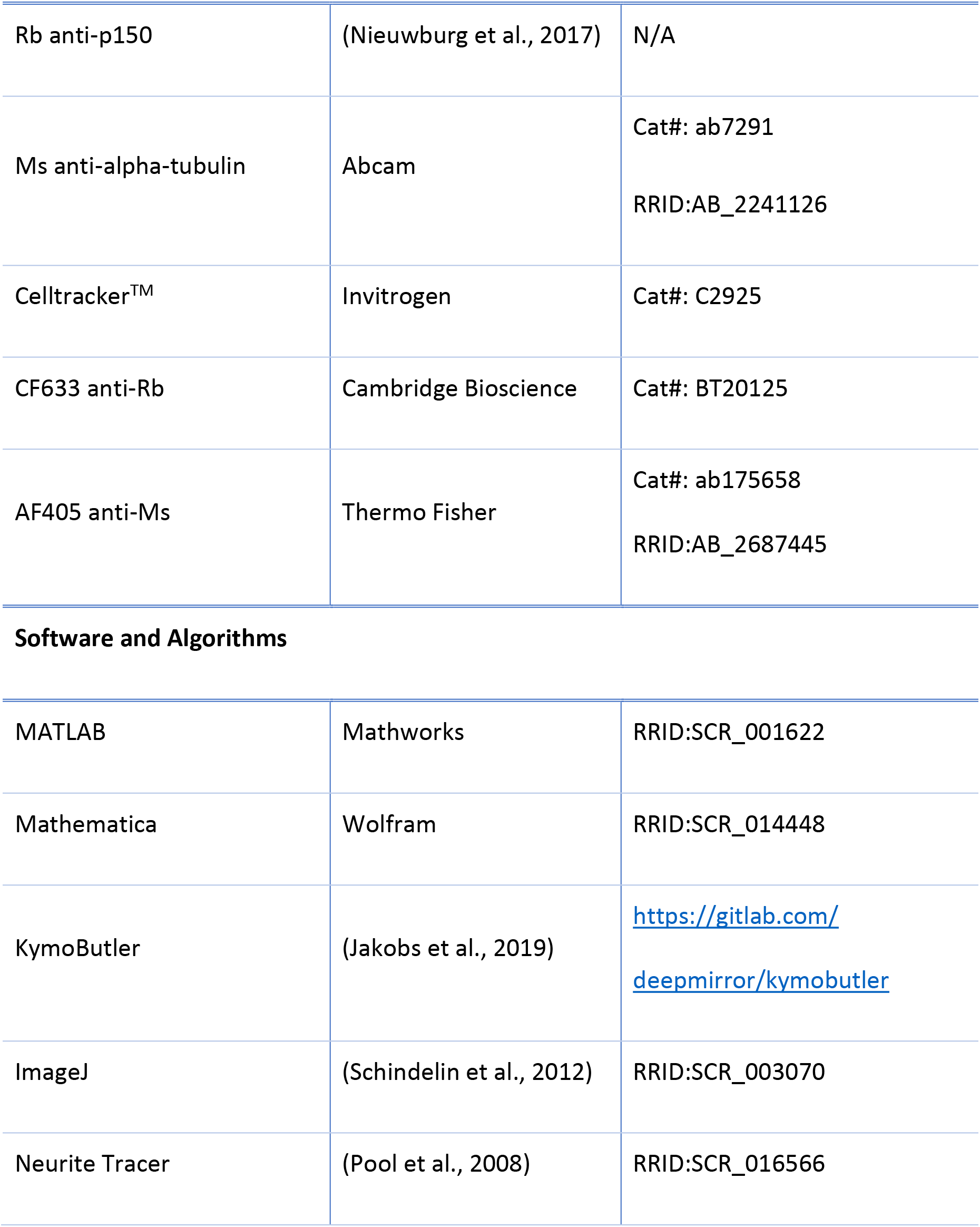

| Reagent or Resource | Source | Identifier |
| --- | --- | --- |
| **Experimental Models:** |  |  |
| **Organisms/Strains** |  |  |
| *D. melanogaster: EB1-GFP* | (Bulgakova et al., 2012) | N/A |
| *D. melanogaster: ubi EB1-GFP* | (Shimada et al., 2006) | N/A |
| *D. melanogaster: Jupiter-mcherry* | (Bergstralh et al., 2015) | N/A |
| *D. melanogaster: p150<sup>1</sup>* | Bloomington | RRID:BDSC_504 |
| *D. melanogaster: p150-RNAi* | Bloomington | RRID:BDSC_3785 |
| *D. melanogaster: khc-RNAi* | Bloomington | RRID:BDSC_35770 |
| *D. melanogaster: elav-gal4* | Bloomington | RRID:BDSC_458 |
| **Antibodies** |  |  |
| Rb anti-p150 | (Nieuwburg et al., 2017) | N/A |
| Ms anti-alpha-tubulin | Abcam | Cat#: ab7291<br>RRID:AB_2241126 |
| Celltracker™ | Invitrogen | Cat#: C2925 |
| CF633 anti-Rb | Cambridge Bioscience | Cat#: BT20125 |
| AF405 anti-Ms | Thermo Fisher | Cat#: ab175658<br>RRID:AB_2687445 |
| Software and Algorithms |  |  |
| MATLAB | Mathworks | RRID:SCR_001622 |
| Mathematica | Wolfram | RRID:SCR_014448 |
| KymoButler | (Jakobs et al., 2019) | https://gitlab.com/deepmirror/kymobutler |
| ImageJ | (Schindelin et al., 2012) | RRID:SCR_003070 |
| Neurite Tracer | (Pool et al., 2008) | RRID:SCR_016566 |

### Fly stocks

MT +end dynamics were visualized with a transgenic fly line expressing EB1-GFP heterozygously under its endogenous promoter (*wh;+;eb1-gfp/tm6b*, gift from the Brown laboratory in Cambridge) (Bulgakova et al., 2013) or a fly expressing EB1-GFP under a *ubiquitin* promotor (*ubi:eb1-gfp;+;+*, gift from the St. Johnston laboratory in Cambridge) (Shimada et al., 2006). Whole MTs were labelled with Jupiter-mcherry (*wh;if/cyo;Jupiter-mcherry*, gift from the St. Johnston laboratory in Cambridge) (Bergstralh et al., 2015). Other stocks used: *p150^1^* (Bloomington # 504), *uas:p150-RNAi* (Vienna Drosophila Stock Center # 3785), *uas:khc-RNAi* (Bloomington # 35770), *khc27* (Bloomington # 67409), khc17 (gift from the St. Johnston laboratory in Cambridge). *uas* constructs were driven by *elav-gal4* (Bloom# 458, *elav* is a neuron specific promotor that ensures the construct is only expressed in the CNS (Yannoni and White, 1997)) and transgenic lines were generated through standard balancer crossing procedures.

### Primary cell culture

3rd instar larvae were picked 5-8 days post fertilisation, and their CNS dissected similarly to (Egger et al., 2013; Sanchez-Soriano et al., 2010). As described in (Egger et al., 2013) the resulting primary culture comprised a mixture of terminally differentiated larvae neurons such as peripheral neurons alongside precursors cells and immature neurons of the adult fly brain. Thereby, the larval CNS lends itself to the study of a heterogenous population of neurons. The CNS tissue was homogenized and dissociated in 100 μl of Dispersion medium (Hank’s Balanced Salt Solution (1xHBSS, Life Technologies, 14170088) supplemented with Phenylthiourea (Sigma-Aldrich P7629, 0.05mg/ml), Dispase (Roche 049404942078001, 4mg/ml), and Collagenase (Worthington Biochem. LS004214, 1mg/ml)) for 5 minutes at 37**°**C. The media was topped up with 200 μl of Cell Culture Medium (Schneider’s Medium, Thermo Fisher 21720024) supplemented with insulin (2 μg/ml Sigma I0516) and fetal bovine serum (1:5 Thermo Fisher Scientific A3160801)) and cells were spun down for 6 min at 650 rcf. The pellet was resuspended in Cell Culture Medium at 5 brains/120 μl. Cells were grown at 26**°**C for 1.5 hours in a droplet of 30 μl Cell Culture Medium in a glass bottom dish between a Concanavalin A-coated glass slide and an uncoated glass slide on top. Initially the cells were cultured with the coated coverslip facing down. After 1.5 hours the chambers were flipped so that cells that did not attach floated off to the opposite (uncoated) side. Culture times were: 4-26 hours (for measuring MT orientation profiles in short and long axons), 22-26 hours (for measuring MT dynamics, both Patronin-YFP and EB1-GFP), and 22-48 hours (for measuring MT dynamics in dendritic processes).

To measure the effect Nocodazole has on MT orientation in axons, the medium was supplemented with 5uM of Nocodazole (Dissolved in DMSO, Sigma-Aldrich M1404-2MG) approximately 12h post-plating and 12h before measuring MT dynamics. The control cells were treated with 0.025% DMSO in culture medium. Treatment and corresponding controls were always run in parallel, and when possible, from the same fly stock. *Uas* driven overexpression was controlled with a fly expressing both *elav::gal4* and *eb1-gfp* to control for the expression of gal4 protein.

To measure the effects of osmolarity changes in the surrounding medium we increased the osmolarity of the culture medium by approximately 100 mOsm (from ~360 mOsm to ~460mOsm, see also https://www.sigmaaldrich.com/GB/en/product/sigma/s9895) by adding 4mg of NaCl to 1ml Culture medium. Cells were first cultured in normal media for 1.5h. Subsequently, the media was removed and replaced with either fresh media (control) or media supplemented with 4g NaCl. Cells were again imaged after 22-26h post plating.

### Live imaging of MTs

All live imaging movies were acquired on a Leica DMi8 inverted microscope with a 63x objective (oil immersion, NA=1.4, Hamamatsu Orca Flash 2.0 camera) and at room temperature (22-25°C). To reduce autofluorescence during imaging, the culture medium was replaced with Live Imaging Solution (Thermo Fisher A14291DJ). Culture media was not replaced for imaging cells in Nocodazole, DMSO, and osmo+ to enable measurement of MT dynamics in the chosen media. For EB1-GFP imaging, an image (exposure time 500 ms) was taken every 2 seconds for 70-150 frames depending on sample bleaching. When imaging both EB1-GFP and Jupiter-mcherry simultaneously, one image was taken every 3 seconds for 100 frames (exposure 500ms). Lamp intensity was set to the lowest level that enabled visual identification of labels.

### p150 antibody staining

24 hours after plating, the cells were treated with 5μM of Celltracker™ (Invitrogen C2925) dye for 30 minutes to label cells in green. Subsequently, cells were fixed in pre-warmed 4% paraformaldehyde (PFA) (pH 7.2, 26°C) for 50 minutes. Post-fixation, the cells were washed in PBS once and then incubated with mouse alpha-tubulin 1:1000 (Abcam ab7291) and rabbit glued/p150 antibody 1:500 (gift from the St Johnston laboratory, (Nieuwburg et al., 2017)) diluted in PBST (2x phosphate-buffered saline (PBS, Oxoid BR0014G) tablets in 400ml H2O + 1.2 ml Triton X-100) + 0.01 g/ml bovine serum albumin at 4°C overnight (~14h). After two quick washes in PBS, the cells were incubated with the secondary antibodies Alexa Fluor 647 (far-red, Thermo Fisher A-21236) and 405 (blue, Thermo Fisher A-31556) for 1.5 hours at room temperature. After another two quick PBS washes, the cells were mounted in Fluoromount (Thermo Fisher 00-4958-02) and imaged.

Images were analysed by drawing a line along axon processes from the base of the axon to its tip in the tubulin channel. The intensity profiles for all 3 channels (p150, tubulin, and Celltracker) were extracted and normalized by their respective median values, and the p150 channel was subsequently divided by the normalised Celltracker channel and the data binned in bins of 10 μm. Finally, the resulting binned *p150-RNAi* p150 profiles were normalized by the mean fluorescence of the respective *wild-type* control p150 fluorescence. The resulting profiles (per axon) were then pooled over all biological replicates and plotted in *Mathematica*. See **Error! Reference source not found.** for a single cell workflow example.

### EB1-GFP dynamics

Kymographs of EB1-GFP tracks in *Drosophila melanogaster* axons were generated by first using the Neurite Tracer plugin in ImageJ to draw lines along axons or dendrites from the centre of the cell body to the farthest EB1-GFP comet signal, i.e. the distalmost growth event (Pool et al., 2008). Subsequently a custom *Mathematica* (https://wolfram.com) algorithm automatically generated kymographs from these lines by plotting the average pixel intensity of 3 adjacent pixels into rows of an image for each frame. The resulting image was then smoothed with a Gaussian kernel of size 3 and wavelet filtered to remove noise. Kymographs were analyzed with *KymoButler* and subsequently post processed in *MATLAB* (https://mathworks.com). Tracks were removed in case: (i) they displaced less than 2 pixels along the x-axis, (ii) they were slower than 1.5 μm/min, (iii) they were faster than 20 μm/min, and (iv) they were visible for less than 4 frames. Additionally, control experiments and their corresponding treatment condition were discarded if the control axons exhibited a mean orientation below 0.8 or average growth velocities below 2 μm/min. To account for outlier comets, the distance from the axon tip was calculated as the distance from the 0.95 quantile EB1-GFP comet.

MT -end polymerisation is much slower than +end polymerisation (Strothman et al., 2019). Since we observed similar growth velocities of cell body-directed and tip-directed MT growth, we are confident that we measured +end out MT growth events in both directions rather than -end growth.

Note that, mature *D. melanogaster* dendrites *in vivo* exhibit a mixed MT orientation (Stone et al., 2008). However, we cultured neurons only up to 48h which might be too short to form fully developed dendrites and our minimal cell culture medium is likely lacking growth factors that would enable further differentiation to form fully -end out dendrites. Additionally, vertebrate dendrites also appear to acquire their characteristic orientation over time (Baas et al., 1989).

### Jupiter-mcherry & EB1-GFP

Kymographs were prepared as for imaging EB1-GFP only (i.e. using Neurite Tracer). Individual shrinkage events were extracted by hand from the resulting kymographs using the ROI tool in *ImageJ* (https://imagej.net). The tracks were then analysed and plotted with *MATLAB* and *Mathematica*. Measuring MT shrinkage dynamics was only possible in regions of low tubulin content, e.g., near the axon tip. We implicitly assumed that MT shrinkage depends neither on MT orientation nor on its position along the axon. However, experimental evidence suggests that a decrease in MT growth length correlates with an increase in shrinkage length (Vasquez et al., 2017), indicating that we likely overestimated MT lengths further away from the axon tip, therefore underestimating the difference between +end out MTs at the tip and those further away from it.

### Statistics

For comparing two groups the Wilcoxon rank sum test was used as implemented in MATLAB (https://www.mathworks.com/help/stats/ranksum.html). The standard error of the mean (s.e.m.) was calculated as 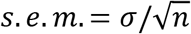. Here *σ* is the standard deviation of the sample and *n* is the number of samples. The 95% confidence interval was calculated by median bootstrapping with 10000 random samples from the distribution. Biological replicates are experiments conducted on different days with different larvae and reagents. We used the Kruskal Wallis test (https://uk.mathworks.com/help/stats/kruskalwallis.html) to compare several samples, followed by a Dunn Sidak post hoc test.

### Solution of the 2-state master equation

We assumed that MTs can either grow or shrink, and each of these two states (*g* and s in short) has a probability distribution that depends on MT length *l* and time *t* ( *p_g_*(*l, t*) and *p_s_*(*l, t*)). MTs can furthermore stop growing and start shrinking with rate *f_g_* = 1/*t_g_* (*t_g_* being the average MT growth time) and stop shrinking to start growing with rate *f_s_* = 1/*t_s_* (*t_s_* being the average MT shrinkage time). Furthermore, MTs are assumed to grow with velocity *v_g_* and shrink with velocity *v_s_*, while they are in the growing- or shrinking-state, respectively. Writing this as a master equation yields:

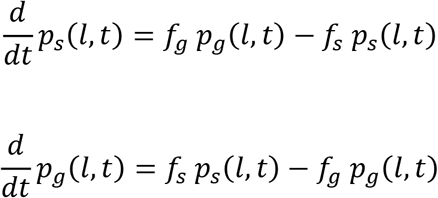

To solve this set of partial differential equations consider the following Fourier transformation of *p_s_*(*l, t*) and *p_g_*(*l, t*):

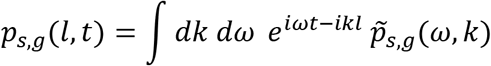

Substituting in the 2-state master equation yields:

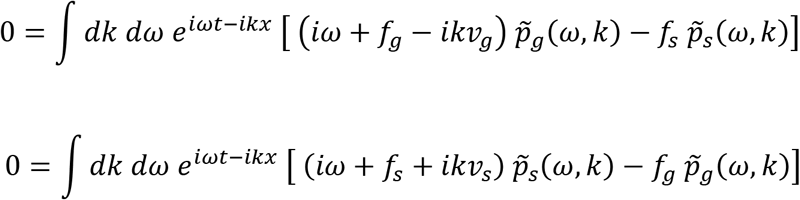

Which can be written as a matrix equation:

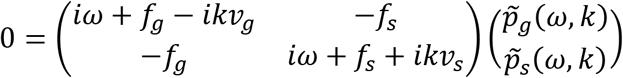

This equation only has non-zero solutions for 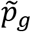 and 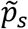 if the matrix determinant is equal to zero:

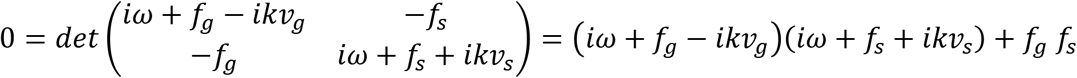

This equation can be written as a dispersion relation:

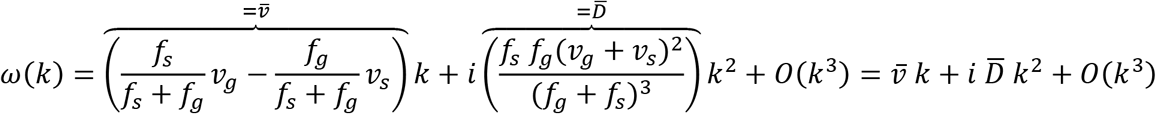

For large times *t* both *ω* and *k* are small so that we can drop terms of the order of *k^3^*. The dispersion relation is then the same as for a diffusion advection process with drift velocity 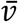 and Diffusion coefficient 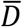. For 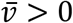 the system will evolve like a diffusion advection process in which MTs would have no average length so that they will become as long as the system allows, i.e. their growth is “unbounded”.

For 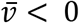, MTs will exhibit an average length that depends on their dynamic parameters which can be calculated as follows: For large times, the overall probability to find a MT with length *l* at time *t* (*p*(*l, t*) = *P_g_*(*l,t*) + *p_s_*(*l,t*)) can be approximated by a modified diffusion-advection equation:

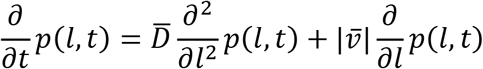

The stationary state 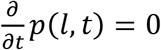 is thus found by:

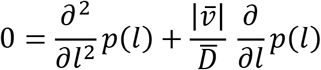

The general solution to this partial differential equation is:

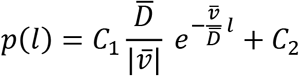

For *p*(*l, t*) to be normalisable: *C_2_* = 0 and 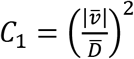. So that:

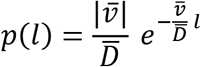

This equation is a 2-state master equation that equates the rate change of a probability to be in one state (i.e. d*p_s_*/dt) to the outflow (-f_s_p_s_, i.e. the likelihood of a shrinking microtubule to start growing) and the inflow (f_g_p_g_, i.e. the likelihood of a growing microtubule to start shrinking) into that state. With *dp/dt* = *∂l/∂t ∂p/∂l* = *v ∂p/∂l* we can write:

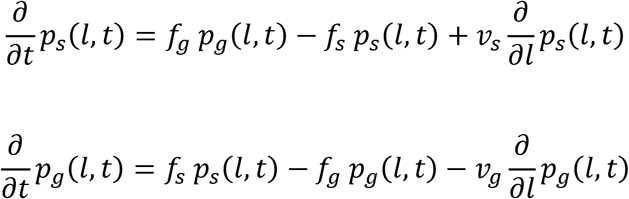

Finally, one can calculate the average MT length *l_MT_* as the expectation value of the length:

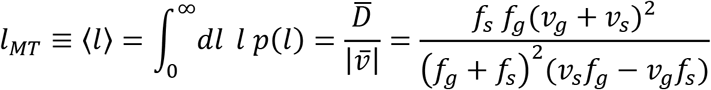

For *v_s_/v_g_* ≈ 1, *f_s_/f_g_* ≈ 1, and *d* = *v/f* the quadratic terms can be Taylor expanded to yield:

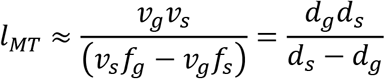

#### Analytical fit to estimate MT growth per cycle based on immunostainings

The p150 fluorescence profiles were calculated as described in the section on “p150 antibody staining”. Subsequently, an exponential function, *p*150(*x*) = *b* + *e*^−*s*(*x*–*x*_0_)^, was fitted to the first 12 bins of the data (corresponds to up until 120 μm from the tip). Here b, s, and x_0_ are fitting parameters with units: [b]=A.U., [s]=1/um, [x_0_]=um. Next, we assumed that *d_g_*(*x*) = *A p*150(*x*)^*α*^, i.e. MT growth per cycle is a simple power law in the p150 fluorescence intensity. *A* and α are fitting parameters and their values are shown in **Error! Reference source not found.**. The expected growth length per cycle for a MT that starts growing at position *x* towards (1) or away (−1) from the cell body can be approximated as:

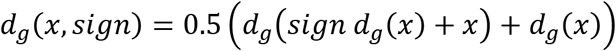

Here we assumed that the average length added to a MT during growth is the mean between the average expected growth length at start and end position of the MT +end. This function was subsequently fitted to the growth length data presented in **Figure 1F.** To do so, the experimental data was first binned in bins of size 10 μm (like the staining data). Then we calculated the integral of *d_g_*(*x, sign*) over each bin for each direction of growth and minimised the squared difference to the experimental results by varying *A* and *α*. Note that we assumed that MTs that grow way from the tip have to be at least 4 μm away from it (average MT length (Yu and Baas, 1994)) and that MTs that grow into the tip may penetrate it by 2 μm.

#### MT sliding simulations

Details of the simulation can be found in (Jakobs et al., 2020). We here present a brief description that focusses on the novel way in which new MTs are added during the simulation. MTs were arranged with their long axis along the *x*-axis of a Cartesian coordinate system and their centers on a hexagonal lattice in the *y* - *z* plane. For simplicity all MTs were assumed to have the same length, l_MT_ = 4μm. The inter-MT spacing in the *y* - *z* plane (~ 30 nm) was assumed to allow individual molecular motors (here, cytoplasmic dynein) to intervene between adjacent filaments and cross-link them with their respective ‘cargo’ or ‘walking’ domains. The simulation was initialized with 10 randomly oriented MTs that were randomly distributed on a hexagonal lattice of lenght 6 μm. New MTs were added to the system depending on the chosen nucleation model:

1. Sliding only: MTs were added at random locations with random orientation every 1100 seconds (~18 minutes). The time was optimised to yield axons of approximately the same length as cultured ones.
2. Sliding and templating: MTs were added at random locations every 1100 seconds. The likelihood of being +end out was calculated by counting the number and orientation of MTs at the location (the center of the MT) in which the MT is added. Then the number of +end out MTs was divided by the total number of MTs to calculate the probability of getting a +end out MT. Finally a random number is drawn between 0 and 1 to determine the orientation of the added MT.
3. Templating Only: Same as above except that the directionality of the movement of molecular motors along the MTs was eliminated. When MT overlaps became occupied with motors, the motors’ gliding direction towards the +end or the −end of the MTs was chosen at random. Note that this is a highly artificial setting that solely removes the sorting effect of microtubule sliding.
4. Sliding and unbounded growth: A random location along the axon was chosen and a random MT orientation (50/50 +end out/-end out) introduced every 435 seconds. As not every MT nucleated in this model, the rate of influx was selected to be higher to enable the same axon growth behaviour. Subsequently, we calculated the likelihood of exhibiting unbounded growth for a MT with the randomly selected orientation and location. To do so, we first calculated the average added length per growth cycle in 10μm bins (distance from the axon tip and separately for +end out and -end out MTs) for each axon in the dataset presented in **Figure 1A-G**. For each bin we then queried whether growth was bounded (added length below 2.2μm) or unbounded. The likelihood of unbounded growth was calculated for each bin by counting the number of axons that exhibited unbounded growth in the bin and dividing that number by all axons. Subsequently, two exponential functions were fitted to the +end out and -end out MT data respectively to determine a function that gives the likelihood of unbounded growth for +end out and -end out MTs as a function of distance from the axon tip. Finally, the random location and the predetermined orientation were used to look up the likelihood of unbounded growth and the MT was assumed to have nucleated successfully when a randomly drawn number [0,1] was smaller than that likelihood.
5. Unbounded growth only: Same as above but molecular motors were again assumed to not have a preferred direction as in 3.
6. Sliding, templating and unbounded growth: A random location along the axon was chosen every 435 seconds and its orientation likelihood calculated as in 2. Subsequently, the unbounded growth likelihood was calculated as in 3. MTs only successfully entered the system if exhibiting unbounded growth.

MTs that were neighbours on the *y-z* plane and overlapping along the x-axis were crosslinked by cytoplasmic dynein. For simplicity and due to the tight packing of MTs in the bundle, only motion in parallel to the *x*-axis was considered. MT velocities were determined by solving a set of force balance equations that characterize dynein interaction with the MTs, detailed in (Jakobs et al., 2020). Furthermore, the left boundary was a leaky spring; MTs that moved into the left boundary were subject to a force of 50 pN/μm and were able to leave the axon with a fixed rate per MT (0.00024/sec). The rate was adjusted to lead to axons of 50 μm in length after approximately 24 hours simulation time. The right boundary was a constant force of 50 pN as described in (Jakobs et al., 2020).

Axons were simulated for 50001 iterations (~28 hours) and all results averaged over 50 separate simulations. Simulation parameters were as follows:

| Symbol | Description | Value | Reasoning |
| --- | --- | --- | --- |
| $\chi$ | Fraction of overlapping<br>MTs that are cross linked | 1 | We previously explored how changing $\chi$<br>affects MT sliding (Jakobs et al., 2020; 2015).<br>In this manuscript we simply wanted to<br>explore the effect of different MT addition models on MT orientation in which we fixed the value at 1. |
| $\lambda$ | Number of motors bound in an overlapping region [#/$\mu\text{m}$] | 5 | We quantified (by eye) the number of MT crosslinks in EM images of axons (Hirokawa et al., 2010) which was approximately 3 per 100 nm. |
| $l_{MT}$ | MT lengths [$\mu\text{m}$] | 4 $\mu\text{m}$ | Average MT lengths in axons measured in (Yu and Baas, 1994) |
| $\xi$ | Drag coefficient of the axoplasm | 1 pN<br>sec /<br>$\mu\text{m}^2$ | Same coefficient used in (Oelz et al., 2018) |
| $f_s$ | Dynein stall force | 1.4 pN | Same coefficient used in (Oelz et al., 2018) |
| $v_0$ | Dynein free velocity | 0.86<br>$\mu\text{m}/\text{sec}$ | Same coefficient used in (Oelz et al., 2018) |
| $dt$ | Simulation timestep per iteration | 2<br>seconds | As we showed previously (Jakobs et al., 2020; 2015), this value is a good choice to ensure smooth movements of MTs during the simulation. |

## Acknowledgements

We would like to thank Eva Pillai, Dennis Bray, Michael Takla, and Kevin Chalut for inspiring discussions and proofreading, Andreas Prokop and Cristina Melero for teaching *Drosophila* dissection techniques, and Sarah Bray, Dmitry Nashchekin, Daniel St Johnston, and Nick Brown for providing *Drosophila* strains, laboratory space and a great atmosphere to work in. The authors acknowledge funding from the Wellcome Trust (PhD studentship 109145/Z/15/Z to MAHJ), the UK Biotechnology and Biological Sciences Research Council (Research Grant BB/N006402/1 to KF), the European Research Council (Consolidator Award 772426 to KF), and the Alexander von Humboldt Foundation (Alexander von Humboldt Professorship to KF).

## Supplementary Information

Figures S1-S9

