## Supplementary Figures for "Unrestrained growth of correctly oriented microtubules instructs axonal microtubule orientation"

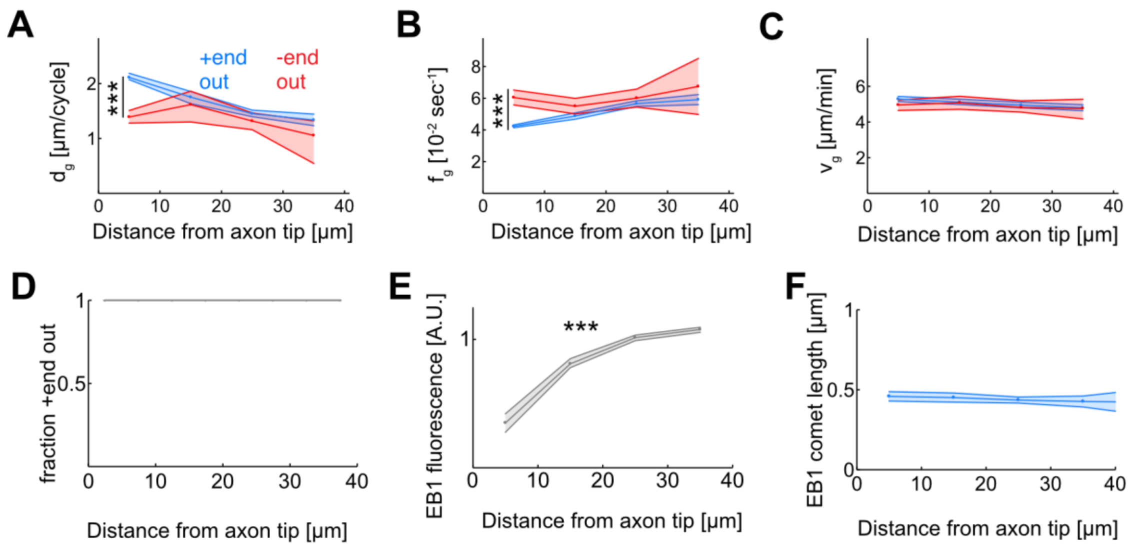


**Supplementary Figure 1: EB1 dynamics as a function of the distance from the axon tip. (A-C)** Microtubule growth. **(A)** Added MT length per growth cycle, $d_g$, **(B)** Catastrophe frequency, $f_g = 1/t_g$, and **(C)** growth velocity, $v_g$, for +end out (blue) and -end out (red) MTs. While MT growth velocities are largely independent of MT orientation and localisation along the axon ($p < 0.003$; Kruskal Wallis test, only bin 1 and 4 of +end out growth velocities are significantly different) **(C)**, +end out MTs near the axon tip undergo catastrophes less frequently than -end out MTs and MTs further away from the tip ($p < 10^{-37}$; Kruskal Wallis test) **(B)**. The resulting longer growth times $t_g$ lead to enhanced growth of these MTs in each growth cycle ($p < 10^{-37}$; Kruskal Wallis test) **(A)**. **(D)** Median MT orientation as a function of distance from the axon tip. Lines represent 30% to 70% quantiles (all 1). No changes are obersved along the axon. **(E)** Median EB1-GFP fluorescence intensity normalised by median fluorescence as a function of distance from the axon tip. EB1 fluorescence decreases towards the axon tip. Lines represent median ± 95% confidence interval. All bins are significantly different and median fluorescence increases away from the axon tip ($p < 10^{-100}$; Kruskal Wallis test, ***: p 0.001). **(F)** Median EB1 comet length as a function of the distance from the axon tip ($p = 0.25$; Kruskal Wallis test) ***: p 0.001. EB1 comets have similar lengths along the axon.

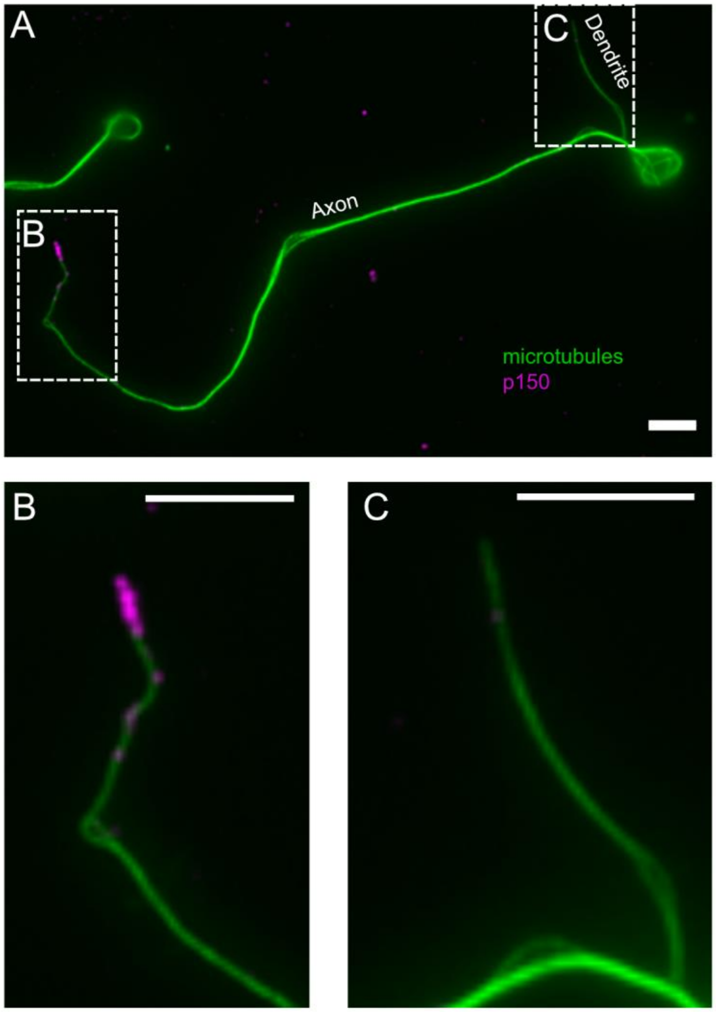


**Supplementary Figure 2: Normalized p150 immunostaining in axonal and dendritic processes.** (A) Representative fluorescence image of a neuron stained for tubulin (green) and normalized p150 (magenta). (B & C) Enlarged images of (B) the axonal tip and (C) the dendritic tip found in the regions defined by the dashed regions in (A). The dendritic process exhibits comparably low levels of p150 if compared to the axonal tip. Scale bars = 5 µm.

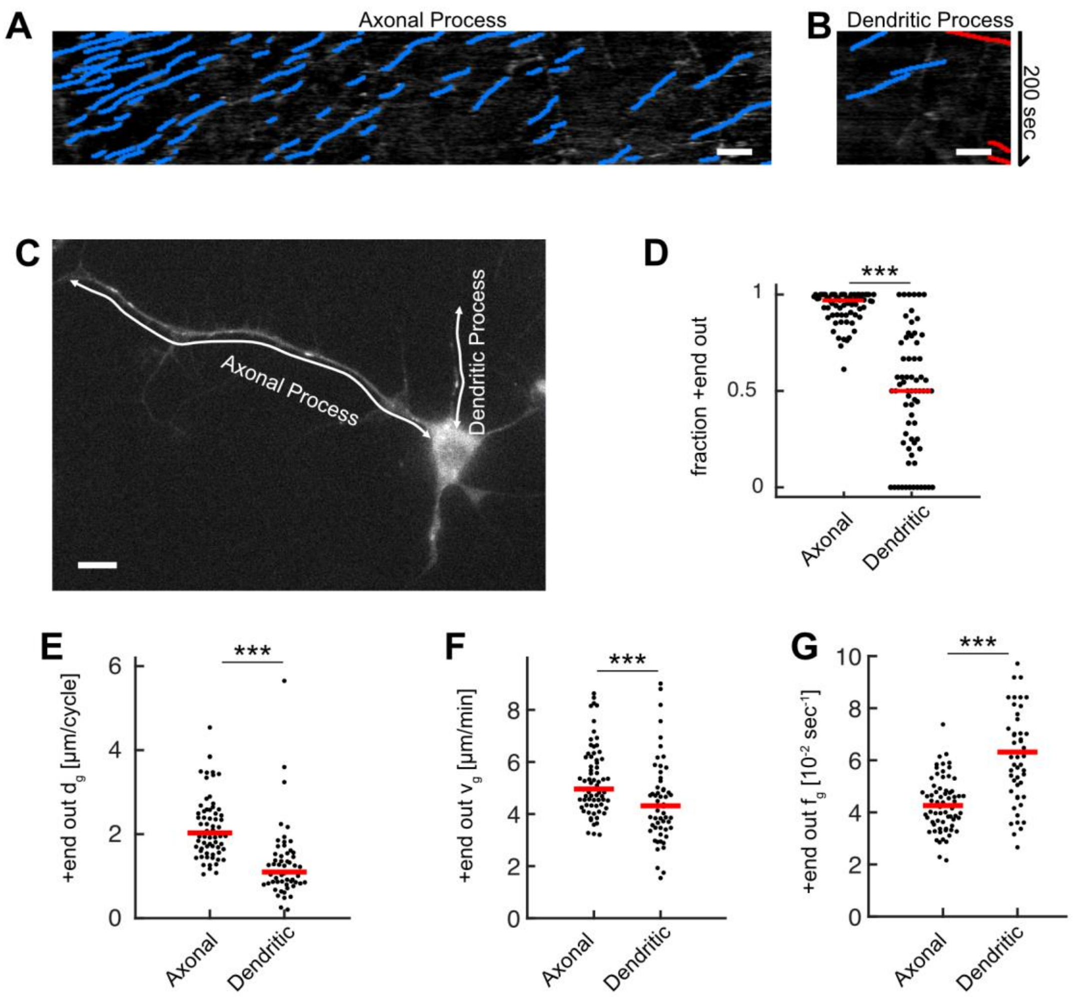


**Supplementary Figure 3: MT growth and orientation is decreased in dendritic processes. (A, B)** Kymographs for **(A)** the axonal and **(B)** the dendritic process shown in **(C)**. **(C)** *D. melanogaster* larva neuron expressing EB1-GFP with an axonal and several dendritic processes. White arrows represent the stationary paths for the kymographs shown in **(A-B)**. **(D)** MT orientation, i.e. the fraction of +end out MTs, for axonal and dendritic processes. Most MTs in axonal but not in dendritic processes have their +ends oriented away from the cell body (N = 69, 20 biological replicates; $p < 10^{-15}$, Wilcoxon ranksum test). **(E)** Growth length per cycle $d_g$ for +end out MTs at the distalmost 10 µm of the axonal or dendritic tips. Black dots: averages for one cell; only cells possessing at least one axonal and one dendritic process (one randomized dendritic process was selected per neuron) were analysed. MTs in axonal processes grew longer than those in dendritic processes ($p < 10^{-8}$, Wilcoxon ranksum test). **(F)** MT growth velocities $v_g$ in the distalmost 10 µm of the axonal or dendritic tips ($p < 10^{-8}$, Wilcoxon ranksum test). (**G)** MT catastrophe frequencies

$f_g$ in the distalmost 10 µm of the axonal or dendritic tips. Catastrophe frequencies are increased in dendritic processes compared to axonal ones ($p < 10^{-8}$ , Wilcoxon ranksum test). Scale bars: 3µm.

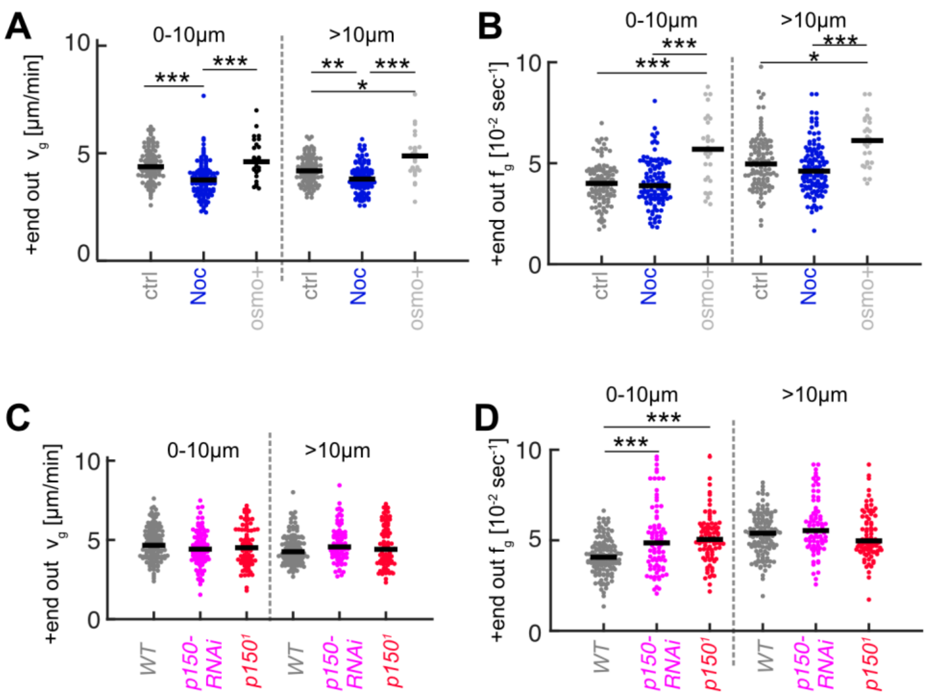


**Supplementary Figure 4: MT growth parameters for Nocodazole and osmo+ treatments (A, B) and p150 knockdown (C, D).** Plots show growth velocities $v_g$ **(A,C)** and catastrophe frequencies $\boldsymbol{f_g}$ **(B,D)** for +end out MTs within 10 µm of the axon tip and further away for the experiments shown in **Error! Reference source not found.** ( *** = p < 0.001, * = p < 0.05, Dunn-Sidak post hoc).

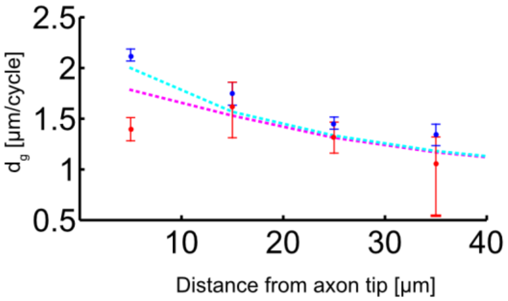


**Supplementary Figure 5: The relationship between p150 protein concentration and the added length per MT growth cycle.** Dots with error bars: experimental data of added length per MT growth cycle $d_g$ as a function of distance from the axon tip is shown in blue (+end out) and red (-end out) (median ± 95% confidence, same plot as in **Supplementary Figure 1A**). Dashed lines: analytic model (see Supplemental methods: 'Analytic model for MT growth as a function of p150 fluorescence' for details), which assumes that $d_g(x)=A$ p150(x)$^{\alpha}$ and $d_g$(+end out, $x$)=0.5($d_g$(-$d_g(x)+x$) + $d_g(x)$) and $d_g$(-end out, $x$)=0.5($d_g(d_g(x)+x)$ + $d_g(x)$). The model thus results in one equation describing the average growth length per cycle, $d_g$ , for +end out MTs, and in another equation describing $d_g$ for -end out MTs. Both equations depend on A, $\alpha$, and the measured p150 fluorescence intensity profile. We simultaneously fitted the predicted $d_g$*'s* for +end out and -end out MTs to the experimentally determined values by varying A and $\alpha$. The resulting pair of parameters was: A = 0.86 and $\alpha$ = 3.62. Our model returned a clear bifurcation between +end out and -end out microtubule growth towards the axon tip, resembling the experimental data and thus suggesting that the observed p150 gradient at the axon tip is strong enough to alter catastrophe frequencies in MTs depending on their orientation.

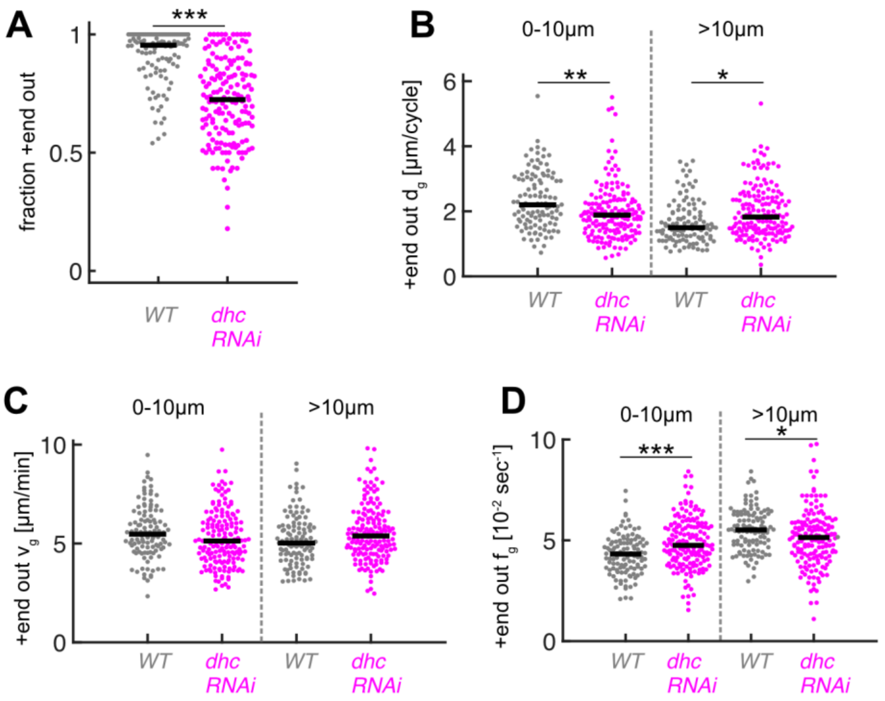


**Supplementary Figure 6: Downregulation of dynein heavy chain expression decreases MT orientation and +end out MT growth at axon tips. (A)** MT orientation (fraction of +end out MTs) for *wild-type* (N=111, 8 biological replicates) and *dhc-RNAi* (N=168, 8 biological replicates). MT orientation was significantly decreased in *dhc-RNAi* axons ($p < 10^{-18}$ , Wilcoxon Rank Sum Test, *** $p < 0.001$). **(B)** +end out MT growth in *wild-type* and *dhc-RNAi*. MT growth in axons expressing *dhc-RNAi* was significantly decreased at the distal-most 10um of the axon compared to control cells and increased further away ($p < 10^{-8}$, Kruskal Wallis Test, ** = $p < 0.01$, * = $p < 0.05$, Dunn-Sidak post hoc). **(C)** Growth velocities $v_g$ of +end out MTs. No significant differences were observed between different MTs. **(D)** Growth time per cycle $t_g$ for +end out MTs. ($p < 10^{-12}$, Kruskal Wallis Test, *** = $p < 0.001$, * = $p < 0.05$, Dunn-Sidak post hoc). Changes in $d_g$ were caused by changes in $f_g$.

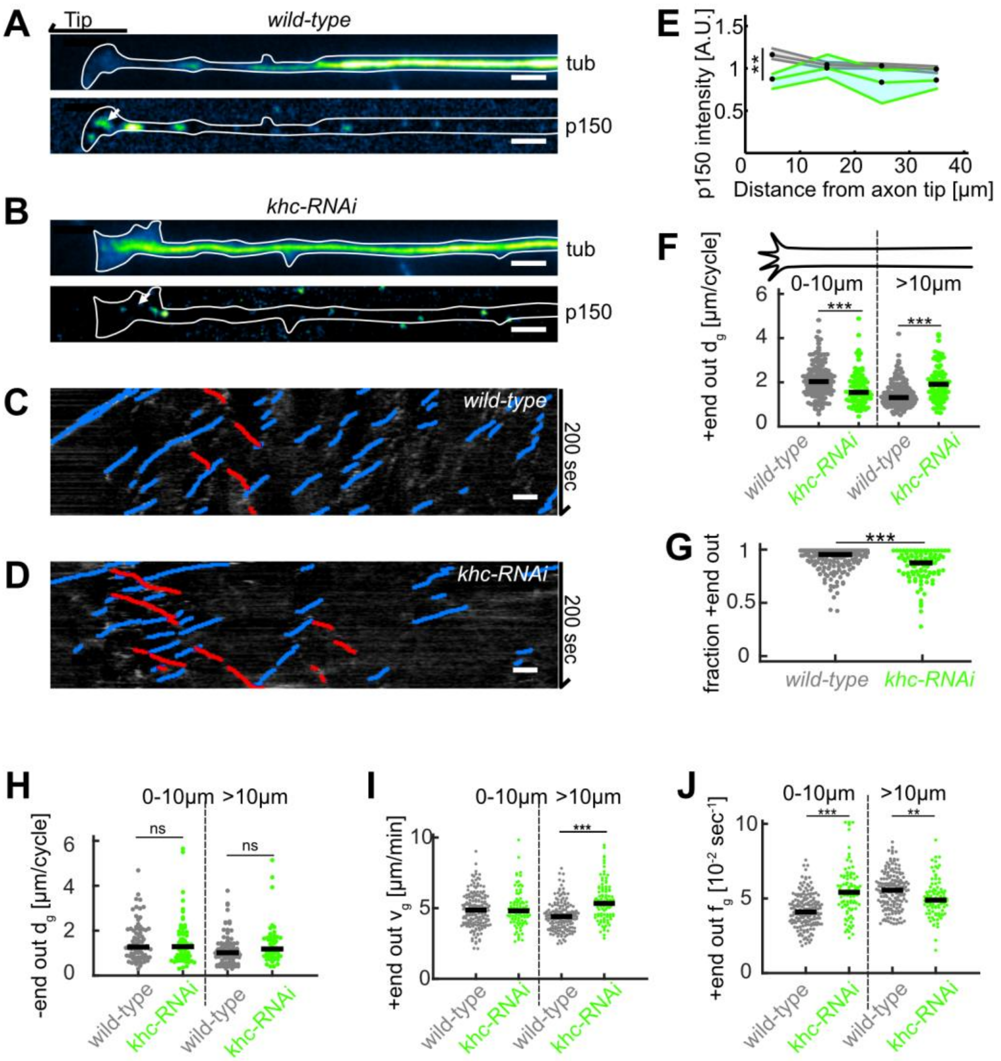


**Supplementary Figure 7: Disruption of kinesin 1 function reduces p150 concentration at axon tips and decreases MT orientation. (A-B)** Tubulin (top) and normalized p150 (bottom) immunostainings of cultured *D. melanogaster* larvae axonal processes. The distal axon tip is pointing left. p150 puncta were more densely clustered around the axon tip in control cells **(A)** compared to axonal processes that expressed UAS driven *khc-RNAi* **(B)**. **(C-D)** Kymographs of EB1-GFP-expressing axons of **(C)** *wild-type* and **(D)** *khc-RNAi*-treated neurons. **(E)** Normalized p150 protein fluorescence as a function of distance from the axon tip for *wild-type* axons (N=162, 5 biological replicates, black line) and *khc-RNAi* axons (N=68, 2 biological replicates, green line). Lines represent median ± 95% confidence interval. p150 protein was less enriched in *khc-RNAi* axon tips ($p < 10^{-16}$, Kruskal Wallis test, 25% downregulation for 0-10 µm: **** $p < 0.01$, Dunn-Sidak post hoc test). **(F)** +end out MT growth in *wild-type* (N=167, 7 biological replicates) and *khc-*

*RNAi* neurons (N=91, 2 biological replicates). MT growth in axons expressing *khc-RNAi* was significantly decreased at the distal-most 10um of the axon compared to control cells. Additionally, +end out MT growth of MTs further away from the tip than 10 µm was increased in *khc-RNAi* compared to control cells ($p < 10^{-9}$, Kruskal Wallis test, *** = $p < 0.001$, Dunn-Sidak post hoc). We can only speculate why the distance grown by MTs further way from the axon tip was increased. It seems rather unlikely that this phenomenon can be explained by impaired dynein-based MT sliding. It is conceivable, however, that the lack of concentrated p150 in the growth cone frees up a pool of tubulin, which is now available for polymerisation further away in the axon. **(G)** MT orientation (fraction of +end out MTs) for *wild-type* and *khc-RNAi* neurons. MT orientation was significantly decreased in *khc-RNAi* axons ($p < 10^{-9}$ , Kruskal Wallis test, $p < 10^{-5}$, *** $p < 0.001$, Dunn-Sidak post hoc test). **(H)** -end out MT lengths per cycle ($p < 0.05$, Kruskal Wallis test). **(I)** +end out MT growth velocities ($p < 10^{-23}$, Kruskal Wallis test, *** = $p < 0.001$, Dunn-Sidak post hoc). **(J)** +end out MT growth times in *wild-type* (N=167, 7 biological replicates) and *khc-RNAi* neurons (N=91, 2 biological replicates) ($p < 10^{-22}$, Kruskal Wallis test, *** = $p < 0.001$, * = $p < 0.05$, Dunn-Sidak post hoc). All scale bars: 2µm.

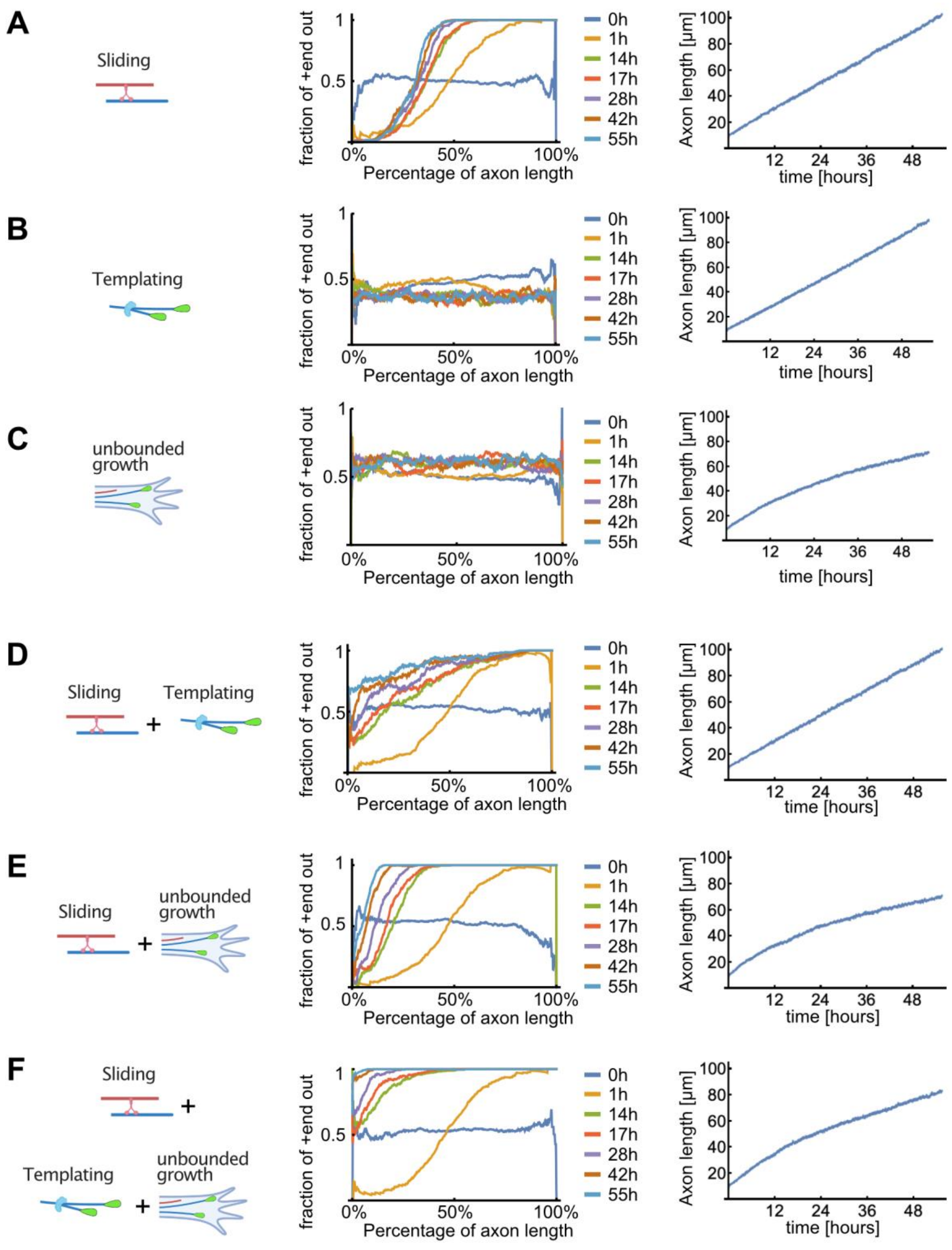


**Supplementary Figure 8: Detailed results of computer simulations for different MT sorting models.** Evolution of the MT orientation profile along the normalized axon length (middle), and of the axon length (right), for different MT sorting models (left). Colours represent simulation time (0-55 hours). **(A)** Dynein-mediated sliding alone. -end out MTs were transported towards the cell body and +end out MTs towards the axon tip. The resulting MT orientation profile was graded along the axon with +end out MTs enriched towards the tip; within approximately 14 hours, the orientation profile became stationary where a fixed fraction of the proximal axon remained enriched with -end out MTs. **(B)** Local bias of MT nucleation (templating) alone. To mimic this situation, the directional movement of dynein motors on the MTs was eliminated by randomly choosing their gliding direction in each MT overlap. As the MT templating mechanism augmented the randomly chosen orientation of the initial MTs in the axon, individually grown axons were either enriched with +end out MTs or with –end out MTs but there was no bias towards either orientation. Consequently, when averaged over many axons, the resulting mean fraction of +end-out MTs was ~50% uniformly. **(C)** Unbounded growth alone. Here a bias towards nucleation of +end-out MTs at the advancing tip of the axon was implemented and calculated from experimental data but motor-induced MT sliding was non-directional as in (B). The resulting MT orientation was stationary along the axon and roughly 55% +end out. **(D)** Dynein-mediated sliding with local bias of MT nucleation (templating). The fraction of +end-out MTs increased along the axon length, and over time, because the biased nucleation resulted in less –end out MTs. However, occasionally in these simulations the dominating MT polarity in the axon was inverted due to 'erroneous' initial nucleation of –end out MTs that then amplified during growth. **(E)** Sliding with unbounded growth of +end out MTs at the tip. The position and orientation of new MTs was chosen at random, but their successful nucleation depended on their likelihood to exhibit unbounded growth as per Error! Reference source not found.. This led to more +end out MTs appearing in the system but not enough to render the axon fully +end out. Furthermore, axon growth ceased to be linear because longer axons were less likely to nucleate new MTs as the region of unbounded growth at the tip did not increase over time. **(F)** Unbounded growth in combination with sliding and templating. New MT positions were sampled from a uniform distribution but with an orientation that depended on the local orientation of nearby MTs. MTs were only added to the simulation if they nucleated as calculated in (C). The fraction of +end out MTs monotonically increased towards the axon tip and the orientation profile gradually become uniform over time, resembling the evolution observed in our experimentally determined profiles in Error! Reference source not found.**B**.

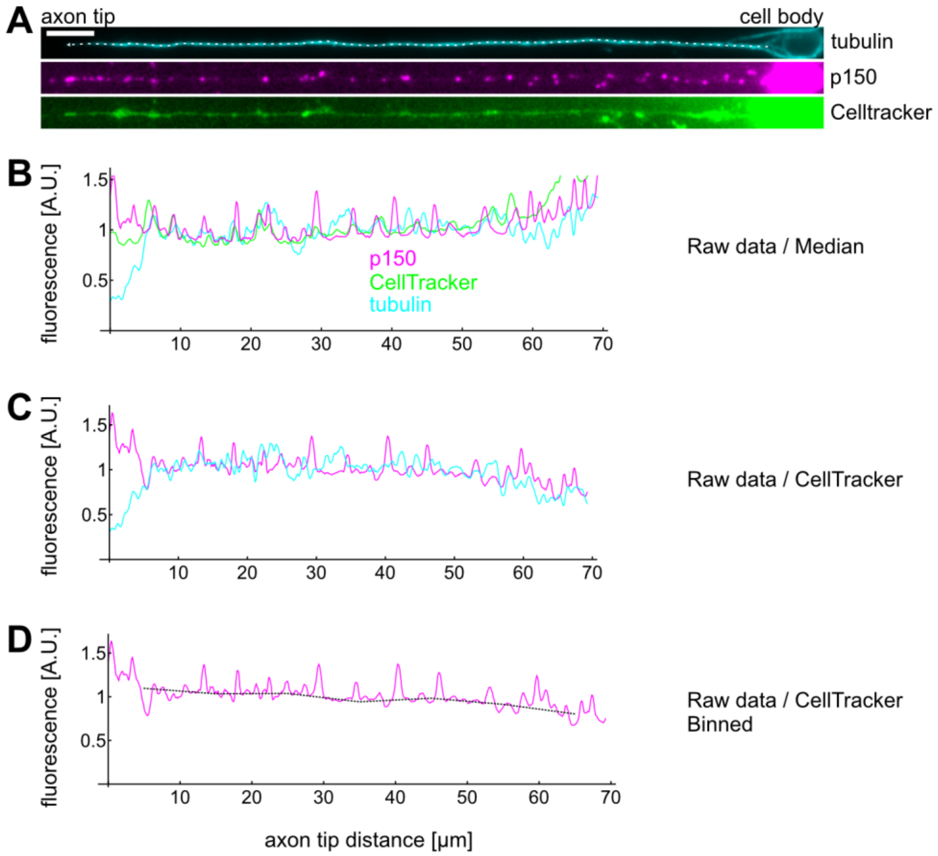


**Supplementary Figure 9: Example calculation of a single axons' p150 profile. (A)** *D. melanogaster* neurons were stained for tubulin, p150, and exposed to CellTracker before fixation. Scale bar: 2 µm. **(A-B)** We then drew a line along the axon from the cell body (right) to the axon tip (left) and extracted intensity profiles for each channel. Typically, we observed a decrease of tubulin and an increase of p150 towards axon tips. Furthermore, all 3 channels exhibited increased fluorescence towards the cell body. **(C)** We then divided the tubulin and p150 profiles by the CellTracker profile to control for increased fluorescence due to larger cell volumes. CellTracker begins fluorescing after permeating the cell membrane and is thereby a good measure for how much cytosol is found at a given location. **(D)** Finally, we binned the data, calculated the mean for each bin, and divided each bin by the average bin value for the corresponding control experiment. The plot in **Error! Reference source not found.C** then shows the bootstrapped median of all bins of all axons of the given condition.
